# Combinatorial microRNA activity is essential for the transition of pluripotent cells from proliferation into dormancy

**DOI:** 10.1101/2023.12.20.572612

**Authors:** Dhanur P. Iyer, Lambert Moyon, Francisca R. Ringeling, Chieh-Yu Cheng, Lars Wittler, Stefan Canzar, Annalisa Marsico, Aydan Bulut-Karslioglu

## Abstract

Dormancy is a key feature of stem cell function in adult tissues as well as embryonic cells in the context of diapause. The establishment of dormancy is an active process that involves extensive transcriptional, epigenetic, and metabolic rewiring^1,2^. How these processes are coordinated to successfully transition cells to the resting dormant state remains unclear. Here we show that microRNA activity, which is otherwise dispensable for pre-implantation development, is essential for the adaptation of early mouse embryos to the dormant state of diapause. In particular, the pluripotent epiblast depends on miRNA activity, the absence of which results in loss of pluripotent cells. Through integration of high-sensitivity small RNA expression profiling of individual embryos and protein expression of miRNA targets with public data of protein-protein interactions, we constructed the miRNA-mediated regulatory network of mouse early embryos specific to diapause. We find that individual miRNAs contribute to the combinatorial regulation by the network and the perturbation of the network compromises embryo survival in diapause. Without miRNAs, nuclear and cytoplasmic bodies show aberrant expression, concurrent with splicing defects. We identified the nutrient-sensitive transcription factor TFE3 as an upstream regulator of diapause-specific miRNAs, linking cytoplasmic mTOR activity to nuclear miRNA biogenesis. Our results place miRNAs as a critical regulatory layer for the molecular rewiring of early embryos to establish dormancy.

## INTRODUCTION

Cells are equipped with a myriad of stress response mechanisms to survive under varying environmental conditions. These confer robustness at the cellular and organismal level and are an evolutionary advantage to species. One such mechanism is embryonic diapause, i.e. the ability to delay post-implantation development by putting early embryos in a dormant state. Embryonic diapause is a widely utilized strategy in over 100 mammalian species, including mice^3,4^. During diapause, development is transiently paused at the blastocyst stage shortly before implantation, with the embryo residing in uterine crypts in close communication with the maternal tissues until receipt of reactivation cues^5^. Under laboratory conditions, diapause can be induced in vivo by surgical removal of ovaries or through hormonal manipulations^6,7^. However, the laborious nature of these experiments and the high number of animals required remain as barriers against identifying molecular mechanisms of this important environmental adaptation.

We have previously shown that blastocysts and embryonic stem cells (ESCs) can be induced to enter a diapause-like dormant state in vitro through direct catalytic inhibition of the prominent growth regulator mTOR^2,8^. mTOR inhibition (mTORi) reduces global anabolic activities in the cell including nascent transcription, translation, and basal metabolic rate^1,8,9^. Mouse blastocysts can be maintained under mTORi for several weeks in vitro^1,8^. Mouse ESCs under mTORi assume a transcriptional and metabolic state highly similar to the in vivo-diapaused epiblast, providing another in vitro system to uncover molecular mechanisms of dormancy entry and exit^8,10^. Both mTORi-paused blastocysts and ESCs maintain developmental competence, as proven by their ability to give rise to live mice or high-grade chimeras, respectively^8^.

Successful entry into dormancy - during diapause and in other systems - requires globally coordinated rewiring of transcriptional, translational, and metabolic activities. Leveraging our in vitro models, we have begun to understand the temporal regulation of dormancy entry during diapause^1,2^. We have shown that immediate targets downstream of mTOR such as translation are inhibited first, followed by chromatin reorganization and a metabolic shift to lipid usage. Still, our understanding of the molecular regulation of dormancy is rudimentary and remains an obstacle to advancing fundamental knowledge about dormancy that may pave the way to improved in vitro technologies.

microRNAs (miRNAs) are small non-coding RNAs that adjust expression levels of target genes by translational or transcriptional interference or mRNA degradation^11^. miRNA-mediated regulation fine-tunes gene expression rather than being the primary determinant of expression levels^11,12^. As such miRNAs are dispensable in many cell types at the steady state. However, miRNA activity is critical for adaptive events like cell state transitions and stress response^13,14^. miRNAs regulate gene expression in an integrative and combinatorial manner, such that multiple miRNAs may target an individual gene and an individual miRNA may target multiple genes. Furthermore, miRNAs often regulate genes related to cell growth, proliferation, and metabolism, which are essential components of the diapause response, making them prime candidate regulators of dormancy. For these reasons, we hypothesized that miRNAs may play a crucial role during the cellular transition from proliferation to dormancy due to their mode of action and target gene groups.

The miRNA Let-7, which is a regulator of developmental timing in C.elegans^15^, has previously been implicated in regulating diapause in the mouse^16^, suggesting that miRNA-mediated regulation of developmental timing may span a wide range of organisms from invertebrates to mammals. Diapause-associated miRNAs have been profiled^17–19^, however, neither the functional significance of implicated miRNAs, nor their role in the larger framework of dormancy has been probed. A broader investigation that integrates miRNAs, their targets, and their upstream regulators is essential to advance our understanding of dormancy and the lack of it currently poses a barrier against determining critical miRNA-regulated processes.

Expression-based methodologies, in conjunction with miRNA-mRNA target prediction algorithms or pre-established miRNA-mRNA target databases, are frequently employed for the purpose of functionally annotating specific miRNAs of interest. These methods facilitate the inference of regulatory relationships between miRNAs and their target genes^20^. While most approaches treat miRNA targets as sets of genes, disregarding the complex gene interactions that they form, network-based approaches can integrate protein-protein interactions (PPIs) and other regulators (e.g. transcription factors) to achieve a systemic view of the miRNA-mediated regulatory mechanisms and identify regulatory hubs and functional modules within the network.

We find here that miRNAs are indispensable for the transition of mouse early embryos and ESCs from proliferation into dormancy in the context of diapause. By ultra-low input sequencing of miRNAs in single embryos, coupled with computational integration of these with proteome profiles, we constructed the miRNA-target network of dormancy and assessed the impact of miRNAs on both direct targets, as well as proteins and pathways downstream of direct interactors. Direct depletion of network-hub miRNAs in embryos reduced the efficiency of dormancy, showing their functional significance. This network adjusts the expression of RNA processing machinery and the organization of membraneless bodies such as PML and P-bodies. Upstream of miRNAs, mining of miRNA regulatory sequences identified the nutrient-sensitive transcription factor TFE3 as part of an mTOR/TFE3/miRNA axis that constitutes a multi-layered program and mediates the transition to dormancy. These results place miRNAs as essential factors in the species’ toolkit to respond to unpredictable conditions during embryonic development.

## RESULTS

### miRNAs are indispensable for the establishment of paused pluripotency

miRNAs are critical regulators of cell state or fate transitions and have been implicated in mediating the response to various stressors^21,22^. We hypothesized that miRNAs may constitute an important regulatory layer in the cellular transition to dormancy in the context of embryonic diapause. We first tested whether miRNAs are functionally required in pluripotent cells during dormancy entry. For this, we induced a diapause-like dormant state in ESCs via mTOR inhibition, as shown before^8^. DGCR8 (also known as PASHA in e.g. C.elegans and D.melanogaster) is a component of the microprocessor complex which produces pre-miRNAs from their primary transcripts^23^. *Dgcr8* KO cells are devoid of mature miRNAs that depend on Dgcr8-dependent canonical processing^24,25^. *Dgcr8* KO early embryos develop and proliferate normally in pre-implantation stages and likewise, DGCR8 is dispensable in ESCs at steady state^25^ (Figure 1A, B), making it possible to identify specific defects associated with the dormancy transition upon mTORi treatment.

**Figure 1.**
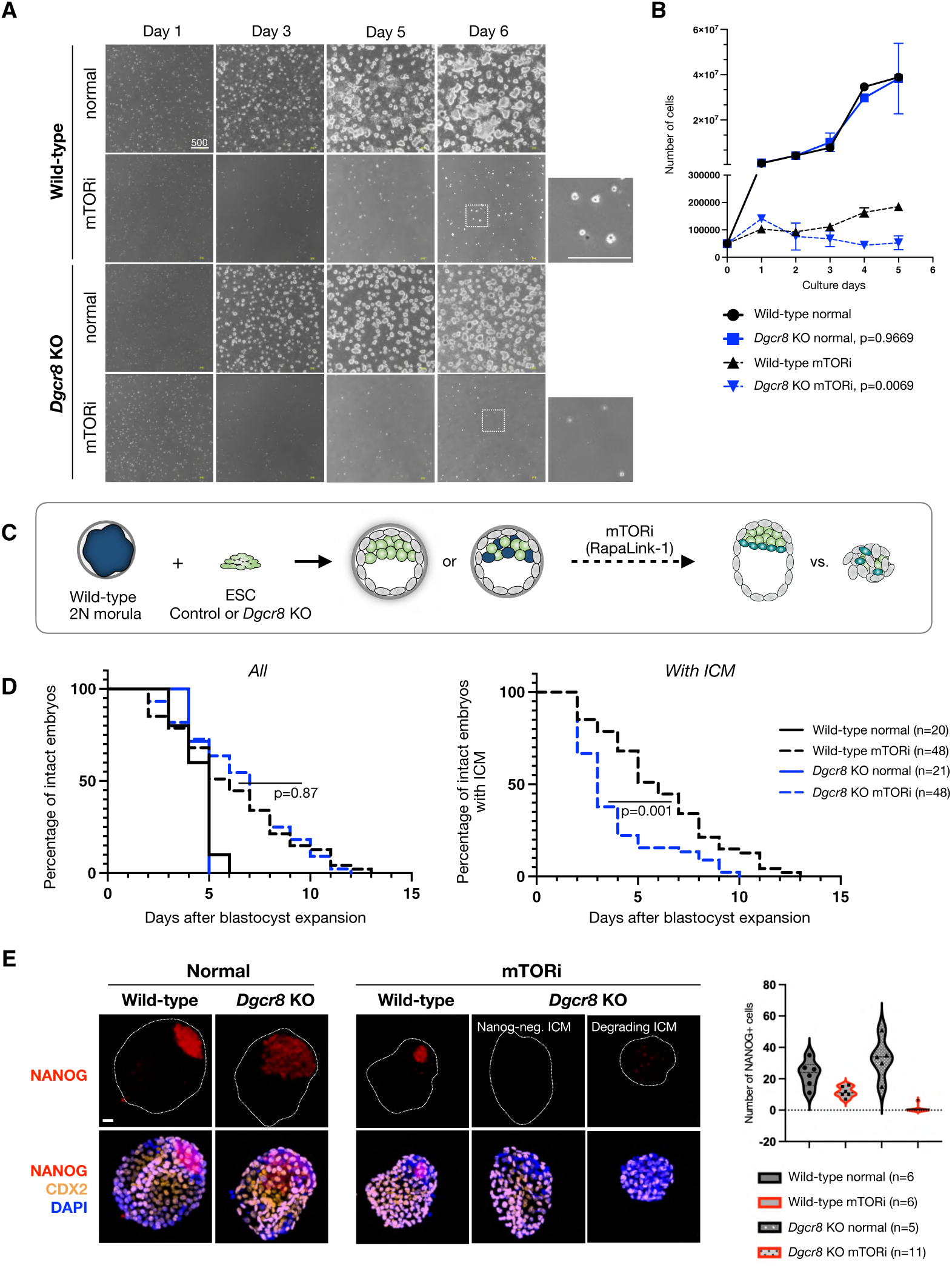
miRNAs are indispensable for the transition of mouse ESCs and embryos into dormancy. **A.** White field images of wild-type and Dgcr8 KO mouse ESCs in normal (proliferative) and dormancy conditions. Cells were induced to enter a diapause-like dormant state via mTOR inhibition as described before8. INK128, a catalytic mTOR inhibitor, was used. Scale bar = 500 um. **B.** Proliferation curves of wild-type and Dgcr8 KO ESCs in normal (proliferative) and dormancy conditions. Cells were plated at low density on 6-well plates and counted on shown days without splitting. Statistical test is nonlinear regression (curve fit), where Dgcr8 KO cells have been compared to wild-type for either normal or mTORi conditions. **C.** Schematics of morula aggregation experiments and the subsequent testing of embryo pausing efficiency. EGFP-labeled wild-type or Dgcr8 KO mouse ESCs were aggregated with wild-type morulae. These were cultured until the blastocyst stage and then treated with DMSO or mTORi. The number of expanded embryos with or without ICM was counted everyday. Embryos with a blastocoel and unfragmented TE were considered intact. Dgcr8 KO ESCs contributed highly to the ICM (Fig. S1). **D.** Survival curves of chimeric wild-type or Dgcr8 KO embryos under mTORi-induced pausing conditions. Left: All intact embryos, right: embryos with a visible ICM. All wild-type embryos retained the ICM during pausing, whereas most Dgcr8 KO embryos lacked it. Statistical test is Mantel-Cox test, with wild-type paused embryos as reference dataset. **E.** Immunofluorescence staining of normal and mTORi-treated (day 3) blastocysts for the epiblast marker NANOG, the trophectoderm marker CDX2, and the DNA stain DAPI. Right panel shows the number of NANOG+ cells in each condition. Between 5 and 11 embryos were stained in each group. The wild-type epiblast reorganizes during pausing in a more compact state, similar to in vivo-diapaused embryos^39^. Dgcr8 KO embryos treated with mTORi either lack the ICM or contain NANOG-cells. Scale bar = 20 µm.

Wild-type and *Dgcr8* KO ESCs proliferate at similar rates under standard ESC culture conditions (serum/LIF, Figure 1A-B). Inhibition of mTOR in wild-type ESCs greatly reduces proliferation while at the same time maintaining morphological features of pluripotent colonies as described before^8^ (Figure 1A-B). In contrast, *Dgcr8* KO ESCs failed to establish dormancy and almost no ESC colonies were visible within 2-3 days of mTOR inhibition (Figure 1A-B). To corroborate this outcome within the setting of embryonic pluripotency, we generated chimeric embryos by aggregating diploid wild-type morulas with either wild-type or *Dgcr8* KO ESCs (Figure 1C). ESCs were labeled with an EGFP reporter randomly integrated into the genome to visualize ESC contribution to the chimeras (Figure 1C, S1A). Both wild-type and *Dgcr8* KO ESC contributed to the inner cell mass (ICM) as expected (Figure S1A). EGFP-carrying cells were more homogeneously present in *Dgcr8* KO ESC-chimeras compared to wild-type, suggesting that *Dgcr8* KO ESCs generate higher-grade chimeras (Figure S1A). After embryos reached the blastocyst stage, they were split into two groups, treated with either DMSO or mTORi, and embryo survival was scored over the next days. Embryos with a blastocoel and without any signs of trophectoderm fragmentation were considered intact. As expected, DMSO-treated embryos did not pause and collapsed within a few days, as these culture conditions are not conducive to implantation or stem cell outgrowth (median survival of 4 days and maximum survival of 5 days after blastocyst expansion, Figure 1D). mTORi-treated wild-type embryos paused development and survived for a maximum of 12 days in culture under these conditions (Figure 1D). *Dgcr8* KO chimeric embryos did not show any difference in overall survival compared to wild-type embryos under mTORi (Figure 1D). However, ∼80% of the *Dgcr8* KO chimeric embryos showed severely compromised ICM within 4 days of mTORi treatment, closely resembling the depletion of *Dgcr8* KO ESCs in culture (Figure 1D-E, S1B). Some embryos had fragmented ICM remnants, while others had an ICM without Nanog expression (Figure 1E, S1B-C). Thus, DGCR8 is essential for the successful transition of ESCs and early embryos into dormancy in the context of mTORi-induced diapause.

### *Dgcr8* KO ESCs show an aberrant transcriptome and differentiation propensity under mTORi-induced dormancy

To get the first insights into how *Dgcr8* KO may affect gene expression and pathway usage during mTORi-induced dormancy transition, we performed bulk RNA-seq in wild-type and *Dgcr8* KO cells in normal ESC culture and after 48h of mTORi treatment (Figure S2A, most *Dgcr8* KO cells are depleted from the culture after this time point). Principal component analysis revealed the genetic background as the primary variable (PC1: 79% variance), and mTORi vs normal conditions as the secondary (PC2: 14% variance) (Figure 2A). Wild-type and *Dgcr8* KO cells showed a similar distribution of significantly differentially expressed (DE) genes, with a higher number of DE genes in KO cells (Figure 2B-C, Table S1, of note: Dgcr8 KO cells have overall lower nascent RNA output compared to wild-type cells, Figure S2B-C). To overcome the strong association of the transcriptomes with the genetic backgrounds, we took a two-step approach to identify the processes that may specifically fail to adapt to mTORi in the KO condition. For this, first, the DE genes (mTORi/normal) were identified in wild-type and KO cells (Figure 2B). Then gene ontology (GO) analysis was performed for the functional allocation of these genes to cellular processes (Figure 2D, Table S2, GO Biological Processes). Afterwards, the GO terms retrieved from wild-type and KO cells were compared, and commonly or distinctly featured terms were identified (the top ten most significant GO terms in each category are shown in Figure 2D). This analysis revealed the aberrant upregulation of genes promoting morphology of differentiation in *Dgrc8* KO cells, in line with the loss of pluripotent cells as seen earlier (Figure 2D, Fig 1E). Downregulated processes were largely shared between wild-type and Dgcr8 KO cells in mTORi and mainly included metabolic terms, suggesting that their regulation may be independent of miRNAs (Figure 2D). These analyses thus further revealed a need for miRNAs for proper transcriptional adaptations as the cells transition from proliferation into dormancy.

**Figure 2.**
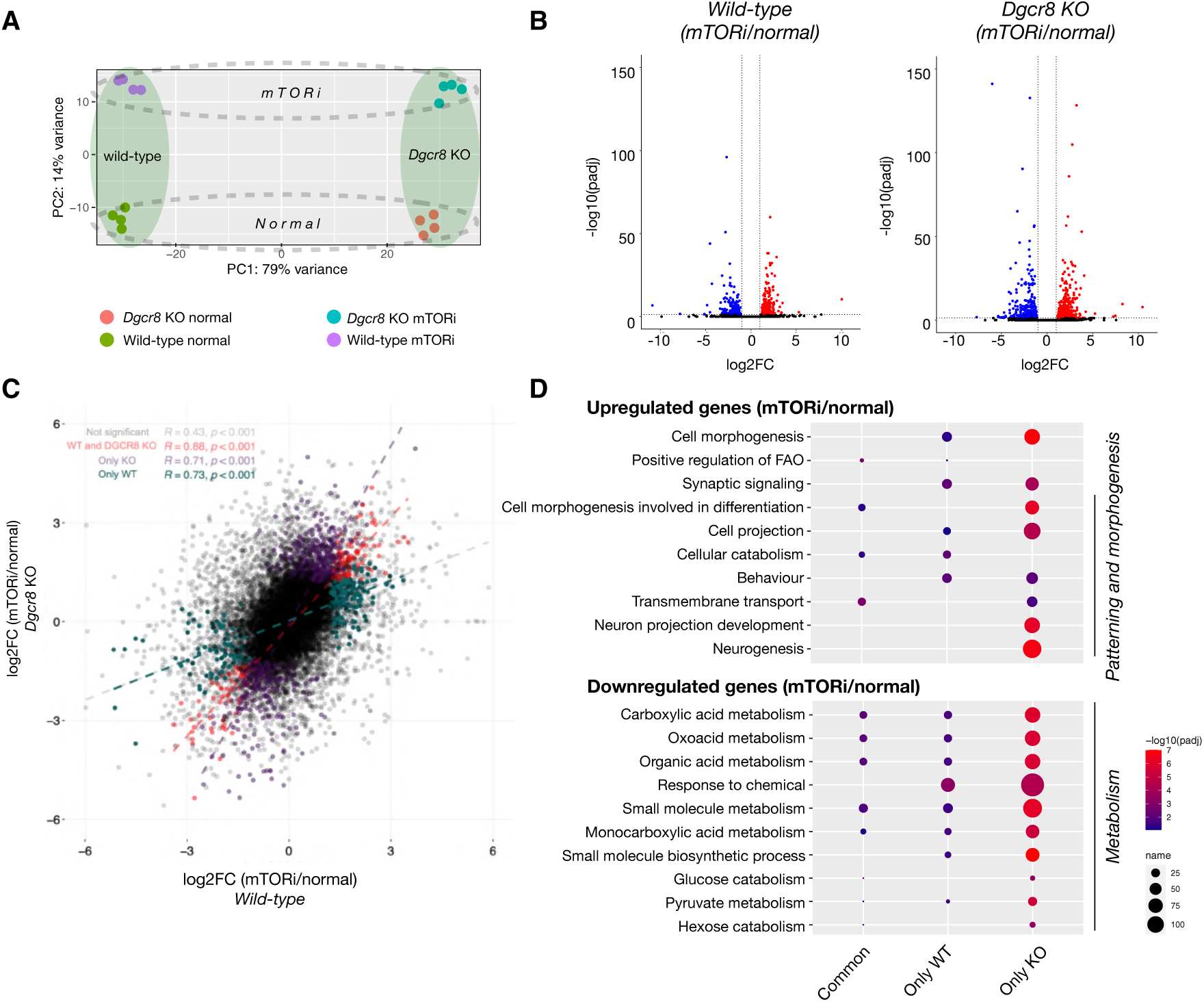
Dgcr8 KO ESCs show an aberrant transcriptome. **A.** PCA plot of wild-type and Dgcr8 KO transcriptomes in normal and mTORi (day 2) conditions. Since most Dgcr8 KO ESCs die after day 3, day 2 was chosen for profiling. **B.** Volcano plots showing differentially expressed genes (padj<0.05, log2FC>1) in wild-type and Dgcr8 KO cells in normal and mTORi conditions. Dgcr8 KO shows more DE genes (738 vs 403 in wild-type, 153 in common) and an overall higher significance level. **C.** Scatter plot showing fold change (mTORi/normal) of each gene in wild-type and Dgcr8 KO cells. Dgcr8 KO cells show a wider log2FC range. DE genes in each background and common DE genes are highlighted with colors. **D.** Gene ontology analyses of significantly downregulated or upregulated genes in wild-type and Dgcr8 KO cells in normal and mTORi conditions. Commonly regulated pathways in both backgrounds are shown together with those that differ based on the genetic background. Top 10 terms are shown, full list is provided in Table S2.

### Single-embryo profiling reveals rewiring of global miRNA expression in dormancy

Embryonic cells fail to enter dormancy in the absence of miRNAs. To begin to understand miRNA-regulated genes and function during this important cellular transition, we first profiled miRNA expression with high sensitivity via ultra-low-input small RNA sequencing (Figure 3A). Diapaused embryos show varied morphologies and survival, therefore miRNA expression profiles may show high variation between individual embryos. For this reason, we generated small RNA expression profiles of individual embryos. Normal late blastocysts (E4.5) were used as control. In vivo-and in vitro-diapaused wild-type embryos were profiled side-by-side with control embryos. In vivo diapause was induced as described before via ovariectomy, followed by progesterone injections^6^. In vitro diapause was induced via mTOR inhibition^8^. miRNAs are tissue-specific regulators of gene expression, thus the different cell types in the blastocyst, especially of embryonic vs. extraembryonic fate, may express distinct miRNAs. miRNAs have a particularly prominent role in the placenta, which we reasoned might confer their progenitor cells, namely trophectoderm (TE), with a specific miRNA signature. However, the TE tissue is not of a uniform state and comprises more naive vs committed cells at the polar and mural ends, respectively^26^. Mural TE shows lower mTOR activity and ceases to proliferate sooner than the polar TE in diapause, attesting to distinct growth dynamics of the mural and polar TE^2,5^. Furthermore, mouse embryos specifically implant from the mural end. For these reasons, we chose to dissect the blastocysts into polar and mural parts and profile small RNAs with higher spatial resolution than individual blastocysts (Figure 3A). Blastocysts were laser dissected to separate the ICM and the adjacent polar TE from the mural TE (Video S1). In addition to blastocysts, we also profiled the ESC and trophoblast stem cells (TSCs) as in vitro derivatives of the respective tissues in the embryo. With this experimental design, we aimed to address three main questions: 1) Do in vivo-and in vitro-diapaused embryos show similar miRNA expression profiles?; 2) Do stem cells and embryos show similar miRNA expression profiles?; and 3) Which miRNAs show consistent expression patterns across conditions?

**Figure 3.**
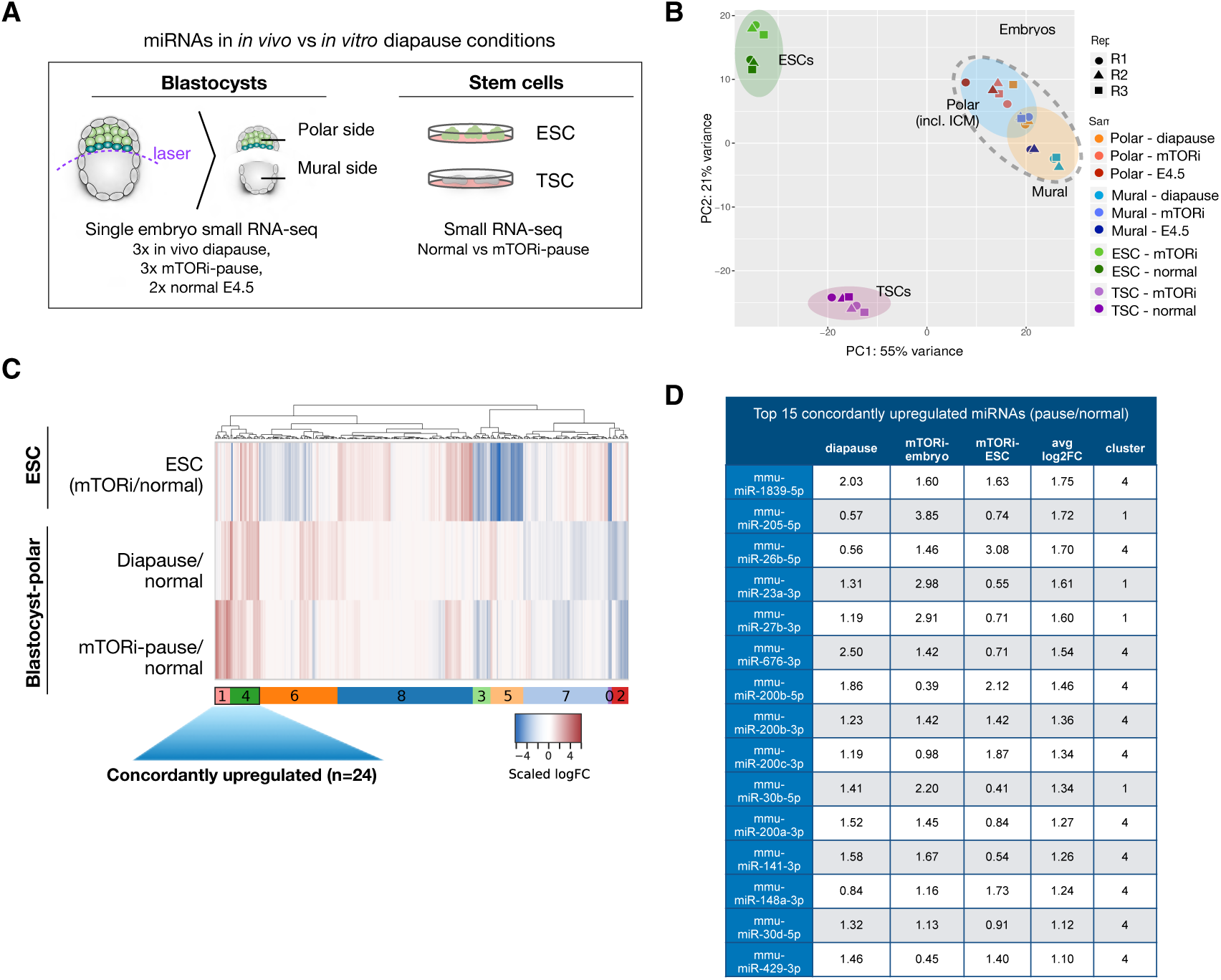
Upregulation of a set of miRNAs is associated with dormancy. **A.** Schematics of sample preparation for small RNA profiling. In vivo diapaused, in vitro diapaused, and normal E4.5 mouse blastocysts were dissected via laser microdissection to separate polar and mural ends (see Video S1). ESCs and TSCs were cultured under standard conditions with or without mTORi. Small RNA expression profiles of single embryo parts and bulk stem cells were generated via low-input small RNA sequencing. **B.** PCA plot of small RNA-seq datasets. Stem cells and embryos are separated along PC1 (55% variance). The polar embryo, which includes pluripotent cells, cluster closer to ESC, whereas the mural embryo clusters closer to TSCs (PC2, 21% variance), suggesting small RNA expression profiles reflective of tissue of origin. In vivo diapaused embryos show higher variability than other groups. **C.** Heatmap showing miRNA expression changes in diapaused embryos (in vitro and in vivo) compared to normal blastocysts, and in paused ESCs compared to normal ESCs. miRNAs were clustered into 9 clusters based on their expression levels in the three samples (see corresponding silhouette in Fig. S3A). Clusters 4 and 1 show a concordant trend of upregulation on average and are the focus of the rest of the study. **D.** Top 15 concordantly upregulated miRNAs and their log2FC values in each condition.

To gauge the agreement of miRNA expression profiles across all samples, we first performed principal component analysis (Figure 3B). Embryonic samples clearly separated from stem cells on PC1 (55% variance), yet ESCs and TSCs were more proximal to their corresponding tissues on PC2 (21% variance, Figure 3B). Within the embryonic cluster, the polar and mural profiles clustered separately. Inter-embryo variability was minimal in the mural part and more pronounced in the polar part, hinting at more dynamic regulation. In vivo diapaused embryos showed higher variability compared to in vitro diapause.

To identify specific patterns of miRNA expression in dormancy, we computed log fold changes (FC) of miRNA expression in paused vs normal embryos and stem cells, and subsequently applied hierarchical clustering to identify groups of miRNAs with concordant or discordant regulation patterns across conditions and tissue types (Figure 3C and S3A, optimal cluster number was identified according to silhouettes in Figure S3B-C, Tables S3-S4). This analysis revealed more similar miRNA expression changes between embryos in diapause (in vivo and in vitro) compared to stem cells, suggesting that tissue complexity or culture media play important roles in miRNA rewiring (Figure 3C, S3A). We saw a greater number of miRNAs in clusters showing a positive increase in expression (mTORi/normal) across polar embryo-ESCs comparisons (two clusters [1 and 4], total of 35 miRNAs) than mural embryo-TSC (Figure S3A, one cluster [number 4], total of 22 miRNAs). In general, we observed very little to no agreement between polar embryo-ESC vs. mural embryo-TSC clusters suggesting distinct regulation in these tissues. Notably, the polar embryo/ESCs showed more dynamic changes compared to mural embryo/TSCs in response to dormancy cues (Figure S3A-S3B) and therefore are the focus of the rest of the study. Amongst the 35 miRNAs in clusters 1 and 4 (Figure 3C), a subset of 24 miRNAs were each concordantly upregulated in all diapause conditions (top 15 are shown in Figure 3D). These are highly enriched for miR-200 family miRNAs such as miR-200a/b/c, miR-141, and miR-429 among others. Taken together, these results reveal dynamic miRNA alterations upon dormancy induction and show that in vitro embryonic diapause largely recapitulates the miRNA regulation of the in vivo phenomenon.

To corroborate the miRNA expression changes in dormancy with an independent approach, we additionally performed miRNA expression analysis in ESCs using the Nanostring technology. The amplification-free sample preparation for Nanostring allows avoiding the artifacts that arise from library preparation for Illumina sequencing. Comparison of miRNA expression profiles of mTORi-treated vs control ESCs generated by RNA-seq and Nanostring showed a positive correlation between the two approaches (Figure S4, Table S5, Spearman coefficient r=0.382, p=2.91e-12). miR-200 family and miR-26b-5p miRNAs were among the top upregulated miRNAs in both assays (Figure S4). This multi-tissue, multi-approach characterization thus revealed a shared regulatory base of in vivo and in vitro diapause, particularly in embryos, and yielded a set of concordantly upregulated miRNAs across diapause conditions.

### Constructing the miRNA-target network of the paused pluripotent state

miRNAs act in a combinatorial fashion, cooperatively fine-tuning many genes at once. We thus conceptualized miRNAs as a regulatory layer that mediates the transition to dormancy, likely via synergistically controlling multiple processes. MiRNAs can exert indirect regulation on functionally associated proteins, influencing the assembly of protein complexes and cellular pathways. To achieve a bird’s eye view of miRNA-regulated processes during dormancy entry, we focused on elucidating the role of miRNAs within the context of protein interaction networks, rather than considering them solely in the context of isolated lists of target genes.

Therefore we constructed an *in silico* post-transcriptional regulatory network of miRNAs, their potential target genes and their interacting proteins (Figure 4A). miRNA-target interactions were retrieved from miRDB and miRTarBase^27,28^ and protein-protein interactions (PPIs) from BioGrid, together with top-scoring interactions from StringDB^29,30^. Only high-confidence interactions from these databases were retained (see Methods) and were exclusively used to connect genes (“nodes”). In addition, we generated deep proteome profiles of mTORi-treated or control ESCs to be integrated with miRNA expression data in the network approach as described below (Figure 4B, Table S6). In order to focus on the most prominent miRNA-mediated changes during dormancy, we have established a scoring framework tailored to prioritize critical edges within the network, specifically concentrating on scoring miRNA-target interactions and Protein-Protein Interactions (PPIs) using two distinct criteria. For miRNA, which play a role in downregulating target genes and thus intricately modulate target gene output, the scoring of miRNA-target edges reflects the expectation that alterations in target gene expression are inversely correlated or discordant with changes in miRNA expression. Namely, miRNA-target edges are scored by subtracting the protein-target log2FC from the connected miRNA log2FC, leading to high-scoring edges when a positively regulated miRNA is connected to a negatively regulated protein. (see Methods). When scoring protein-protein interaction edges, our emphasis lies on the co-expression of the interacting proteins. This approach is rooted in our expectation that perturbations caused by miRNAs ripple through protein complexes and cellular pathways. Furthermore, proteins belonging to the same PPI subnetwork tend to exhibit co-regulation, according to the ‘guilt-by-association’ principle^31^. To transform proteomic data into scores that reflect biological interactions, protein-protein edges were scored by summing the proteomics expression changes of connected proteins (see Methods), leading to high positive or negative scores when two proteins show joint positive or negative regulation. Finally, by keeping the highest scoring edges according to the criteria outlined above (see Methods), we were able to extract connections of highest concordance (protein-protein) or discordance (miRNA-target) in terms of expression changes during pausing. Of note, downregulated miRNAs were not included in the network construction, since our goal was to identify miRNAs that are specifically induced at the time of diapause entry and their putative target genes. From an initial network of 24 upregulated miRNAs and 4,346 proteins, we filtered for the top 10% of miRNA-target edges and the top 1% of PPIs, thus extracting a network comprising 335 PPIs and 196 miRNA-target connections (Fig 4C, non-DE nodes with a single interaction were left out for ease of visualization, the complete network is provided in Table S7).

**Figure 4.**
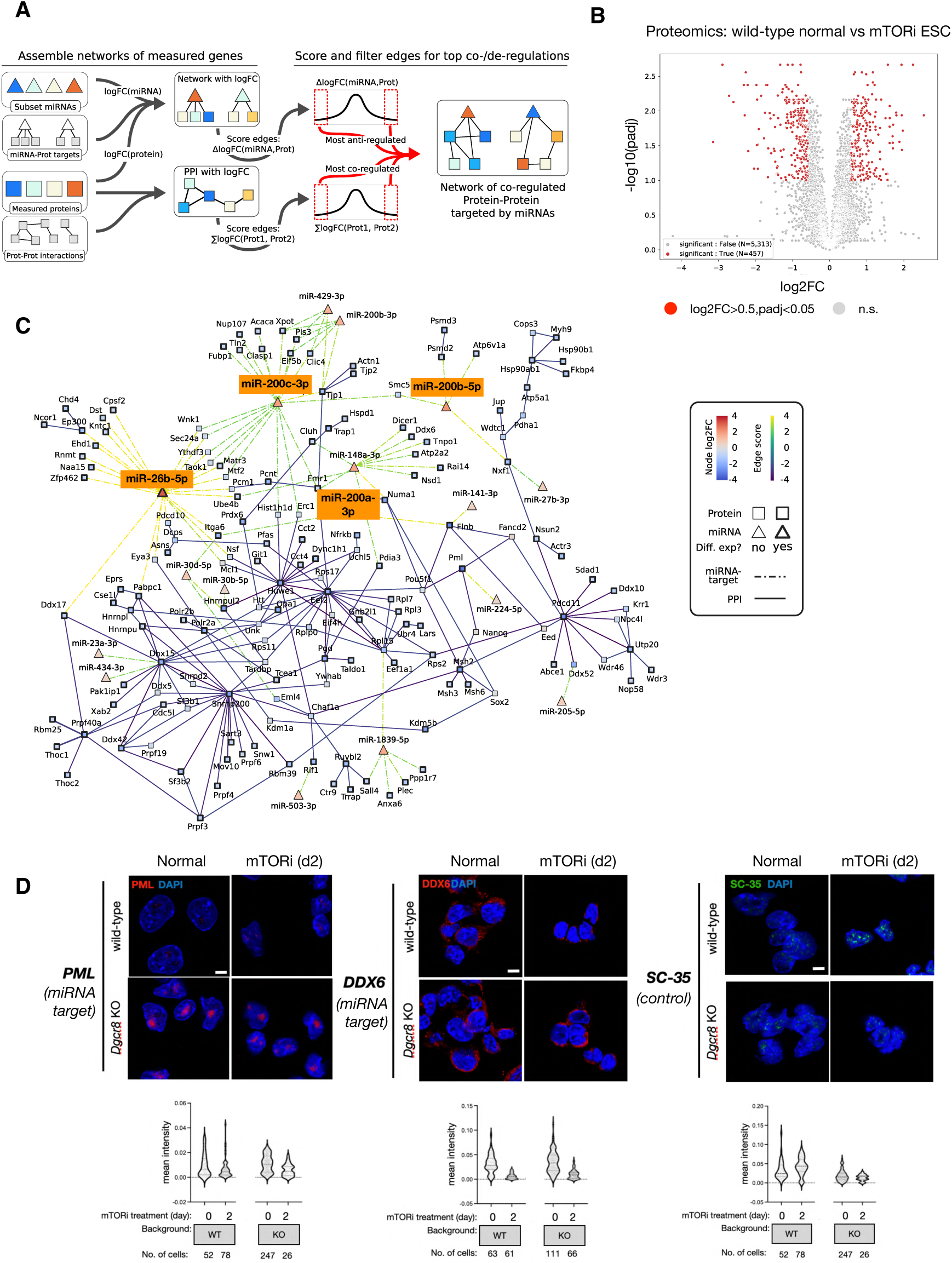
The miRNA-target network of the paused pluripotent state. **A.** Schematics of the computational analysis to construct the network. From the measured logFC of proteins and miRNAs, scores are assigned to edges to assess concordant (for proteins interactions) or discordant regulation (miRNA-target targets). Comparison of actual scores to a distribution from random pairings enables the identification of the most extreme edge scores to keep in the final network. **B.** Differential expression analysis of normal and paused wild-type ESC proteomes. 457 proteins are significantly differentially expressed (216 up, 241 down) using the criteria log2FC>0.5 and padj<0.1. **C.** miRNA network of concordantly upregulated miRNAs and their significantly downregulated target proteins. A stringency cutoff is applied to keep the top 10% of miRNA-target edges, and top 1% of protein-protein interactions. P-value = 1.327e-30 from Fisher exact test (one-sided) for enrichment in differentially expressed protein. A less stringent network with the first cutoff lowered to the top 30% of miRNA-target edges is in Figure S5B, while an ESC-only network is provided in Figure S6. **D.** IF stainings and single-cell quantifications of PML and DDX6 (in-network) and SC35 (control) proteins in wild-type and *Dgcr8* KO cells in normal and pausing conditions (day 2).

The network is enriched in significantly differentially downregulated proteins (119 out of the 431 retained proteins, one-way Fisher’s Exact test p-value = 1.327e-9), while also capturing the majority (17 out of 24) of miRNAs upregulated during pausing in both ESCs and embryos. miR-200 family miRNAs emerged as prominent candidate regulators also within the network, with miR-200b-5p and miR-200c-3p ranking as second and third highest scoring miRNA-target and miR-200c-3p appearing as a prominent “hub”, i.e. exhibiting a high number of regulatory interactions modulating target gene protein levels. Other miR-200 family miRNAs miR-200a-3p, miR-141-3p, and miR-429-3p are also part of the network. The hub node miR-26b-5p is the most upregulated miRNA in the network and PML is the most downregulated protein (Figure 4C). Additionally, including PPIs allows one to visualize how regulatory relationships between miRNA and direct target genes propagate and potentially affect cellular pathways. For example, our results suggest that miRNA deregulation may lead to downstream effects on regulators of RNA processing (mainly pre-mRNA splicing), such as Snrnp200 and Pdcd11, which emerge as prominent hubs in the dormancy network despite not being directly targeted by miRNAs.

While a cutoff of 10% on the scores of miRNA-target interactions allows us to focus on the core network of dormancy (as in Figure 4C), showing the strongest effects on miRNAs and their gene targets; additional, lower-scoring regulatory interactions might contribute to dormancy-related cellular processes. We therefore zoomed out to lower the score cutoff and extend the analysis to the top 30% miRNA-target edges (Figure S5A-B). This revealed other putative regulators of dormancy, including members of the miRNA let-7 family (let-7d and let-7g), which have been previously reported to get upregulated during mouse diapause^16^. The upregulation of let-7 and downregulation of its target genes are however less pronounced than other candidate miRNAs identified in the stringent core network, therefore we exclude it from downstream analyses.

Finally, we explored in detail the ESC-specific regulatory network, independent of the concordance of miRNA expression in embryo samples (Figure S6). Here we wanted to test whether the reduced complexity of the ESC model and the availability of both miRNA expression and matched proteomics data in this cell type would allow identification of candidate miRNAs that remain inconspicuous in the main network. Via this approach, an additional set of 44 miRNAs and 108 new gene targets were found. Interestingly, many of the added miRNAs were found in higher proportion in high-scoring interactions with proteins from the stringent network. For example, PML was additionally targeted by miR-370-5p, miR-92b-5p, miR-1193-3p, miR-291a-5p, and miR-92a-2-5p, amongst the top 25 highest scoring connections. Notably, miR-92, which is an insulin-dependent regulator of L1 diapause in C.elegans^32^, was found upregulated in paused ESCs and is part of the ESC network (Figure S6), suggesting a prominent role in pluripotent cells.

In summary, we present here a core miRNA-target network associated with diapause (Figure 4C). This network is constructed using a combination of miRNA and protein expression data, target prediction, and stringent filtering criteria, retaining only high-scoring relationships characterized by substantial absolute expression changes between miRNAs and their targets, as well as co-expression of functionally related proteins through PPIs. This network provides a foundation for unveiling potential regulatory mechanisms both upstream and downstream of miRNA control during the diapause process.

The in-network proteins were strongly enriched for RNA processing, chromosome organization, and cellular development (Figure S7A). Among these, major organizers of membraneless cellular bodies such as PML (of PML bodies) and DDX6 (of P-bodies) were found (Figure 4C). To test whether these are indeed downregulated at the protein level in dormancy, we treated wild-type and *Dgcr8* KO ESCs with mTORi and performed immunofluorescence (IF) on day 2 of the transition (Figure 4D). As control, we stained for SC-35, which is a major regulator of another membraneless body, the nuclear speckle. SC-35 is not in the miRNA-target network, while PML and DDX6 levels are predicted to be regulated by miRNAs. Single-cell signal quantifications showed that the in-network proteins PML and DDX6 were downregulated in wild-type ESCs upon mTORi, whereas SC-35 was upregulated (Figure 4D). *Dgcr8* KO ESCs in normal ESC culture had higher PML and DDX6 expression compared to wild-type ESCs, whereas mean SC-35 expression was lower in *Dgcr8* KO. Importantly, even though *Dgcr8* KO ESCs did downregulate PML and DDX6 during dormancy entry, overall the silencing was not as efficient as in wild-type ESCs (Figure 4D). In comparison, SC-35 mean levels did not change during mTORi treatment in *Dgcr8* KO cells. These results confirmed that the downregulation of in-network proteins during dormancy entry in wild-type cells and revealed that the expression of these proteins was perturbed in *Dgcr8* KO cells under both normal and mTORi conditions.

### Network perturbation by manipulation of individual nodes leads to loss of pluripotent cells during dormancy transition

Having identified the network of miRNAs that are associated with the diapause entry in embryos as well as ESCs, we next probed the functional requirement for these miRNAs at this cellular transition. We selected the miR-200 family and miR-26b for their high upregulation levels and prominent connectivity within the network (Figure 4B). The requirement for these miRNAs during diapause entry was directly tested in embryos via loss-of-function perturbations. For this, 2 cell-stage embryos were injected with inhibitors against the miR-200 family (pan inhibitor that targets all miR-200 members), miR-200 family together with miR26b-5p, or control inhibitors (Figure 5A, S8A, inhibitors are sequence-specific antimiRs, see Methods). Injected embryos were cultured until the blastocyst stage, and were afterwards treated with the mTOR inhibitor RapaLink-1 to induce diapause in vitro. The percentage of intact blastocysts was scored every day and selected embryos were stained for the epiblast marker NANOG and the trophectoderm marker CDX2 (Figure 5B, C). Inhibition of miR-200 and miR-26 activity significantly reduced embryo survival compared to the control (Figure 5B). Most embryos were lost within the first 5 days of diapause, corroborating the need for miRNAs during this cell state transition. MiRNA-inhibited embryos had fewer NANOG+ cells compared to control embryos, suggesting the disruption of transcriptional networks (Figure 5C). These results show that perturbation of the network reduces the efficiency of in vitro diapause.

**Figure 5.**
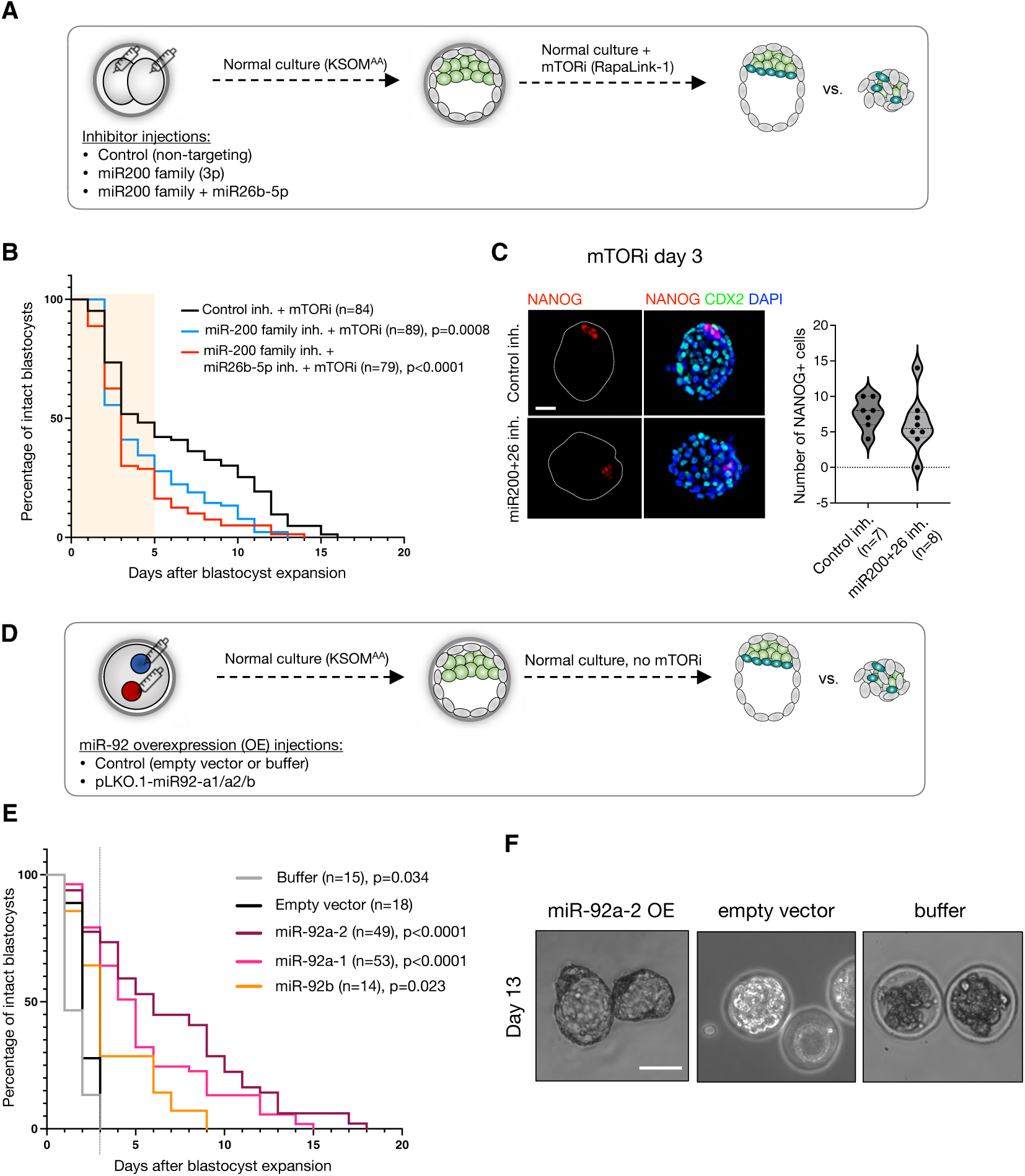
Combinatorial miRNA activity promotes efficient transition to dormancy during in vitro diapause. **A.** Schematics of the experiment. 2 cell-stage embryos were microinjected with synthetic miRNA inhibitors (antimiRs) against the miR-200 family and miR-26b-5p, or a control inhibitor (non-targeting). After injection, the embryos were cultured until the blastocyst stage under standard conditions. Afterwards, the blastocysts were treated with DMSO or RapaLink-1 to induce in vitro diapause. **B.** Survival curves of embryos generated as described above. The number of expanded embryos was counted every day. Embryos with a blastocoel and unfragmented TE were considered intact. Statistical test is Mantel-Cox test with control inhibitor+mTORi as the reference dataset. n, number of embryos. **C.** Immunofluorescence staining of control-injected or antimiR-injected representative embryos on day 3 of mTORi treatment for the epiblast marker NANOG, the trophectoderm marker CDX2, and the DNA stain DAPI. Right panels show the number of NANOG+ cells per embryo in each condition. n= 7-8 embryos were used. Statistical test is unpaired t test with Welch’s correction. Scale bar, 20 µm. **D.** Schematics of miR-92 overexpression (OE). Zygotic pronuclei were injected with the linearized miR92 overexpression construct (pLKO.1 background, U6 promoter), empty vector, or injection buffer. The embryos were cultured in standard medium without mTORi until the end of the assay. For scoring, the same procedure as in (A) is applied. **E.** Survival curves embryos miR-92 OE or control embryos. n, number of embryos in each group. Statistical test is Mantel-Cox test with empty vector injection as the control dataset. Bright field images of miR-92 OE or control embryos on day 13 of in vitro culture without mTORi. Scale bar, 100 µm.

If miRNAs are necessary for the entry into diapause, can they act in a stand-alone manner to directly induce it? To investigate this question, we tested whether miRNA overexpression can bypass mTOR inhibition and induce a diapause-like embryonic state in vitro. For this, we chose the miRNA miR-92 as our top candidate since its parental clusters miR-17-92 and miR-106-363 are highly expressed in pluripotent cells, upregulated in paused ESCs, and its orthologue miR-235 is associated with diapause in C.elegans^32^. miR-92 is transcribed from three distinct loci in the mouse genome on chromosomes 14, X, and 3, giving rise to miR-92a-1, 92a-2, and 92b. To stably overexpress miR-92, parental transcripts of each gene were cloned and integrated into the genome via injection of linearized plasmids into embryos (Figure 5D). Overexpression and processing of the hairpin into mature miR-92a-3p products was confirmed via Taqman qPCR approach (Figure S8B). Once the embryos reached the blastocyst stage, they were further cultured in base media without mTOR inhibitor and survival of intact blastocysts was scored (Figure 5E). miR-92 overexpression was sufficient to prolong the survival of intact blastocysts up to 18 days in vitro as opposed to empty vector or buffer controls (Figure 5E, F). miR-92a-1 and 92a-2, encoded by the miR-17/92 and 106a/363 clusters respectively, yielded higher efficiency compared to miR-92b. Together, these results implicate miRNAs as functional regulators of dormancy in the context of embryonic diapause.

### The cellular transition into dormancy is associated with alternative splicing events

Many cellular stressors result in alternative splicing (AS), which in turn impacts proteome diversity. AS has been linked to mTOR/insulin pathway activity and its infidelity is associated with aging^33,34^. Since the in-network proteins are strongly enriched for RNA processing factors, including RNA splicing, we next asked whether the cellular transition into dormancy is associated with AS (Figure S7A). Indeed, *Dgcr8* KO cells failed to replicate the AS events observed in wild-type cells during dormancy entry (Figure S7B). Regulators of RNA biosynthesis were predicted to be particularly affected in *Dgcr8* KO cells (Figure 7C). We validated 9 AS events via selective amplification of isoforms (Figure S7D). Among these, several genes showed different AS levels in wild-type and KO cells (MBD2C, UBR2, WIPI2; Figure S7D-E). Altered isoform usage level was also validated at the protein level for the MBD2 protein (Figure S7F). Overall, these results reveal altered AS events downstream of mTOR inhibition and their perturbation in cells devoid of DGCR8-dependent miRNA activity.

### The mTOR-TFE3 axis regulates miRNA biogenesis in dormancy

We have so far identified the miRNA-target network specific to embryonic diapause and corroborated its functional significance. Yet, how these miRNAs are selectively upregulated under diapause conditions is unclear. To answer this question and bridge the gap between upstream (cytoplasmic, mTOR) and downstream (nuclear, miRNA) regulators of dormancy, we mined promoters of concordantly upregulated miRNAs for enriched transcription factor (TF) binding sites (TFBS) (Figure 6A). Transcription start sites of miRNAs were retrieved from the FANTOM5 database^35^, and promoters were defined as the region 1,500 bp upstream and 500 bp downstream of the TSS. After intersection with the clustered miRNAs, we predicted the presence of TFBS motifs within the set of the 14 promoters of the concordant up-regulated miRNAs (cluster 1 and cluster 4, Figure 3C). Enrichment of a given motif was evaluated by comparing the fraction of promoters with a TFBS for these candidate promoters versus a background set composed of the promoters of all other detected miRNAs (N=91).

**Figure 6.**
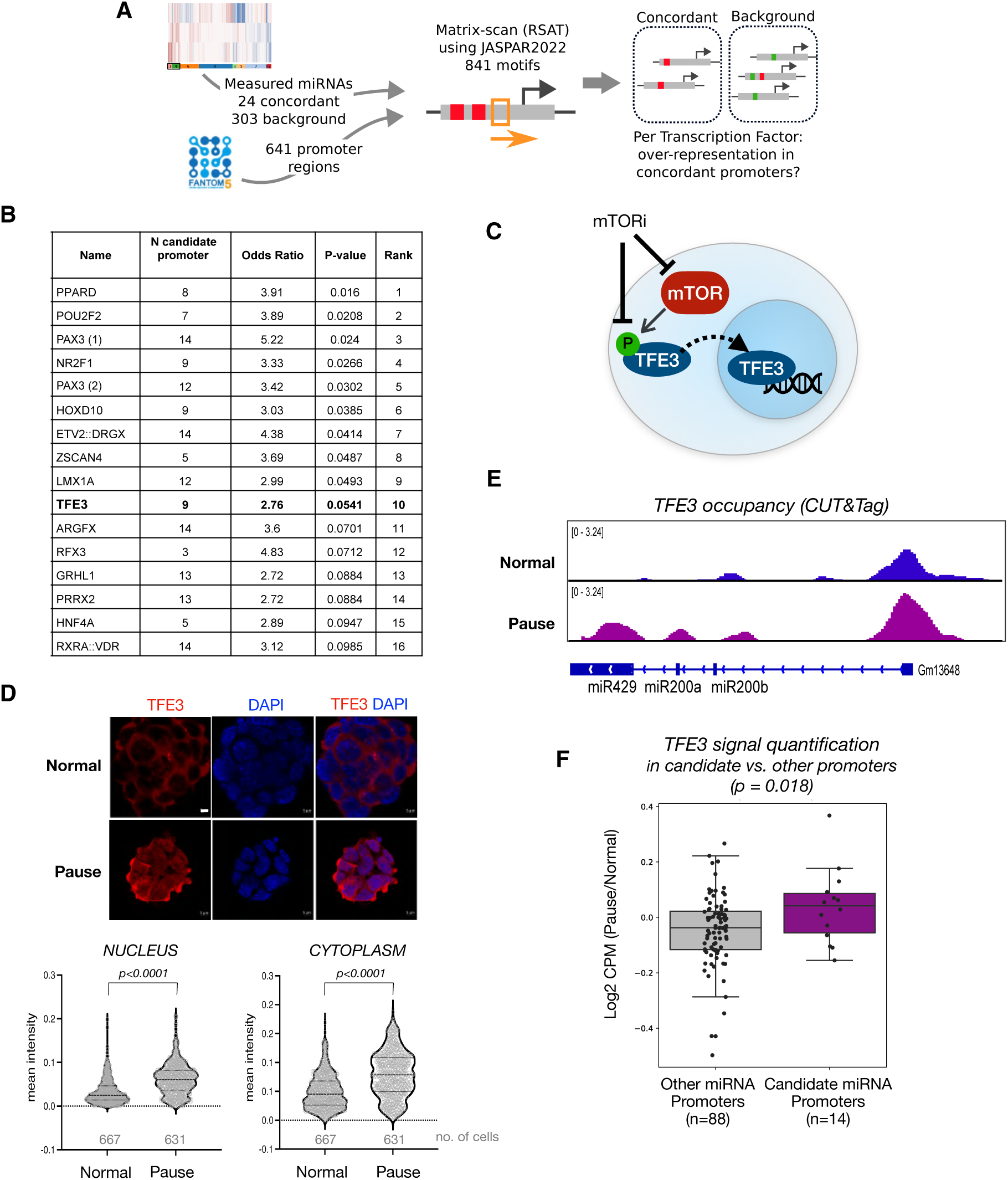
The mTOR-TFE3 axis regulates miRNA biogenesis in dormancy. **A.** Schematics of TF mining at candidate miRNA promoters. Transcription start sites of miRNAs were retrieved from the FANTOM5 database and used to scan the [-1,500, +500] regions with JASPAR motifs. High-confidence hits were then used to compare the fractions of promoters with a given motif, for promoters of concordant, positive logFoldChange miRNAs against promoters of other measured miRNAs. from concordant up-regulated miRNAs. **B.** Enriched TFs at candidate miRNA promoters. **C.** Schematics of the mTOR-TFE3 axis. mTOR phosphorylates TFE3 when active, which results in its sequestration in the cytoplasm. When mTOR is inactive, nonphosphorylated TFE3 instead translocates into the nucleus for regulation of target genes. **D.** TFE3 staining in wild-type normal and paused ESCs. Lower panel shows single-cell quantifications of mean fluorescence intensity in the nucleus and cytoplasm. Scale bar = 10 µm. Statistical test is Kolmogorov-Smirnov test, two-sided. **E.** Genome browser view of TFE3 occupancy over the Gm13648 gene which contains the miR-200a, miR-200b, and miR-429 miRNAs. TFE3 occupancy was mapped via CUT&Tag in wild-type normal and paused ESCs. **F.** Comparison of the log2 ratio of TFE3 levels (maximum scaled CPM) at candidate (n=14) and control (n=88) miRNA promoters. Statistical test is Mann-Whitney U-test, one-sided.

Via this approach, we identified 15 TFs with significant motif enrichment at candidate upregulated miRNA promoters (Figure 6B). Identified TFs comprise of important developmental regulators (e.g. HOXD10, ZSCAN4, PAX3) and TFs responsive to external stimuli, such as hormones (e.g. NR2F1, RXRA, PPARD) and nutrients (TFE3). To test whether candidate TFs indeed bind candidate diapause-associated miRNAs, we eliminated those that are not expressed in ESCs (e.g. HOXD10, PAX3) and focussed our efforts on TFE3. TFE3 and the closely related factor TFEB are known to regulate ESC metabolism and lysosomal genes via their TF activity within the nucleus^36^. Intracellular TFE3/TFEB localization dynamically changes in an mTOR-dependent manner. When mTOR is active, TFE3/TFEB are sequestered in the cytoplasm, whereas nutrient depletion leads to loss of mTOR-dependent TFE phosphorylation and its translocation into the nucleus^37^ (Figure 6C). We reasoned that nuclear TFE3 activity may induce transcription of target miRNAs in diapause, thus providing the missing link between mTOR activity and miRNA upregulation.

To test this hypothesis, we first visualized TFE3 localization in normal and paused wild-type ESCs (Figure 6D). mTOR inhibition increased overall TFE3 expression in ESCs, including its levels in the nucleus and the same was observed for TFEB (Figure 6D, S9). We have previously shown that mTOR complex 1 activity, which regulates TFE3 phosphorylation, is effectively inhibited by mTORi under these conditions, therefore additional regulators may contribute to TFE3 cytoplasmic localization in paused ESCs. To test whether nuclear TFE3 binds target miRNA promoters in paused ESCs, we mapped its occupancy at chromatin via CUT&Tag (Figure 6E-F). TFE3 indeed bound target miRNA promoters in paused cells at higher levels compared to normal ESCs and compared to control miRNAs (Figure 6F). miR-200a/b, which we functionally validated in previous experiments, were enriched in TFE3 binding and among its top targets (Figure 6E). In summary, we identify an mTOR-TFE3-miRNA axis, which regulates rewiring of miRNA expression and downstream protein levels in the transition of pluripotent cells from proliferation to dormancy.

## DISCUSSION

miRNAs are important regulators of cellular transitions, mediating prompt target downregulation via translational or transcriptional interference or mRNA decay. The prominent role of miRNAs in regulation of growth and proliferation led us to hypothesize that they may regulate diapause entry downstream of mTOR inhibition. Here we show that miRNAs are indispensable for diapause entry by rigorous testing of *Dgcr8* KO ESCs and embryos. By integrating single-embryo miRNA expression profiles with target predictions and proteome datasets, we constructed a miRNA-target interaction network of dormancy entry in the context of mouse diapause. Functional testing of the identified miRNAs directly on embryos further corroborated the necessity of this network for efficient entry of embryos into diapause. We further identified TFE3 as a nutrient-sensitive TF that regulates miRNA biogenesis downstream of mTOR activity. Our results therefore provide the first comprehensive overview of miRNA function in early mouse embryos during the transition into diapause.

Our finding of the essentiality of miRNAs for dormancy entry does not align with their dispensability for mouse pre-implantation development^25, 38^. However, this discrepancy is not without precedent, as other critical stem cell regulators that are otherwise dispensable for preimplantation development such as LIF/gp130 and WNT have been shown to be specifically required in diapause^39, 40^. Accumulating knowledge points to diapause as a distinct pluripotent state that necessitates additional regulation, including stress-response mechanisms, and we identify here miRNAs as such an essential regulatory layer. As our knowledge about diapause-specific dormancy mechanisms expand, it will be important in the future to synthesize an understanding of the mechanisms pertaining to stress response and mediate the cell state transition vs. the mechanisms pertaining to maintaining cellular identity during periods of dormancy. This knowledge may then be transferable to other tissues, mainly in the adult organism, that rely on dormancy-reactivation cycles for regeneration.

miRNAs have been previously associated with the dormant state in other contexts such as larval diapause in non-mammalian species^41–43^. Yet, most efforts are directed towards dissecting individual miRNA-target interactions or general miRNA profiling. As such, we only have a limited understanding of miRNAs as a regulatory layer, including the interconnectivity of miRNAs and their targets. Here we aimed to work towards a larger context of miRNA activity in the cellular transition to diapause. As such, we provide the experimentally verified stringent miRNA-target interaction network and refrain from going into further mechanistic details of miRNA-mRNA binding site biology. Further experiments, such as AGO pull-down and site-specific mutations, will deliver insights into each of the nodes and edges in our network. In addition to our spatially resolved miRNA mapping, single cell-level assays may reveal whether these regulators are required in the pluripotent epiblast, primitive endoderm, or the polar TE - the three cell types that makes up the ‘embryonic’ side of the blastocyst.

miRNAs can be of embryonic or maternal origin. Extracellular vesicles (EV) carrying miRNAs are a medium of maternal-embryo communication during diapause^16^. This can lead to differences in the miRNA expression profiles of in vivo and in vitro diapaused embryos. As we see a large agreement between the two in their miRNA expression profiles, it appears that maternal miRNAs do not globally shift miRNA abundance in vivo. Yet, the maternal contribution cannot be ruled out and warrants further investigation.

The miR200/ZEB1 axis regulates epithelial-to-mesenchymal transition (EMT) and has been extensively studied in cancers^44, 45^. Although diapaused embryos show pre-implantation morphology and gene expression signatures devoid of EMT, it has been shown that the epiblast polarizes in a WNT-dependent manner. Therefore, it is plausible that miR-200 activity may be associated with morphological changes of the epiblast. Several other miRNAs in our network regulate cell cycle progression, including miR-26b and miR-148^46–48^. Thus in addition to the downregulation of PML and DDX6 shown in this manuscript, in-network miRNAs are likely to contribute to dormancy entry via other events such as suppression of cell cycle and cellular growth. In line with this, PCNA (regulator of cell proliferation) and RIF1 (regulator of replication timing) are downregulated, among others, in our network. Depletion of DDX6 has previously been shown to confer ‘hyper-pluripotency’ to ESCs and thus may safeguard pluripotency during diapause^49^.

Uncovering miRNAs as a regulatory layer also offers a prospect for their use in diagnosis. Circulating miRNAs, including mir-200 family and miR-92, have long been proposed for use as cancer biomarkers^50–52^. The use of miRNAs as diapause biomarkers could provide a non-invasive way of diapause detection, particularly in wild species in which detection of diapause via hormonal measures is impractical or unreliable^53^.

## ACKNOWLEDGMENTS

We thank Vera van der Weijden, Igor Ulitsky, Jan-Wilhelm Kornfeld, and Volker Busskamp for discussions and feedback, and the scientific service facilities of the Max Planck Institute for Molecular Genetics for excellent service. We thank Constance Ciaudo for sharing *Dgcr8* KO ESCs and Jennifer Shay, Cordula Mancini, and Birgit Romberg for assistance. This project was supported by the German Academic Exchange Service (DAAD) PhD Fellowship to DPI (91730547), the Deutsche Forschungsgemeinschaft (SFB/TR-84 TP C01) to A.M. and L.M, the Helmholtz Center (AM), the Max Planck Society (AB-K), and the Sofja Kovalevskaja Award (Humboldt Foundation) to AB-K.

## AUTHOR CONTRIBUTIONS

AB-K and DPI conceived and developed the project. DPI performed all experiments except embryo laser dissections, aggregations, and CUT&Tag. Embryo manipulations were performed by LW and the Transgenic Facility of the Max Planck Institute. C-YC performed CUT&Tag, LM performed all computational analyses except bulk RNA-seq and alternative splicing. The latter was performed by FRR under the supervision of SC. AM and AB-K supervised the project. All authors contributed to the manuscript.

## SUPPLEMENTARY INFORMATION

Code available at: https://github.com/marsico-lab/mirna_embryo_pausing

### SUPPLEMENTARY FIGURES

**Figure S1.**
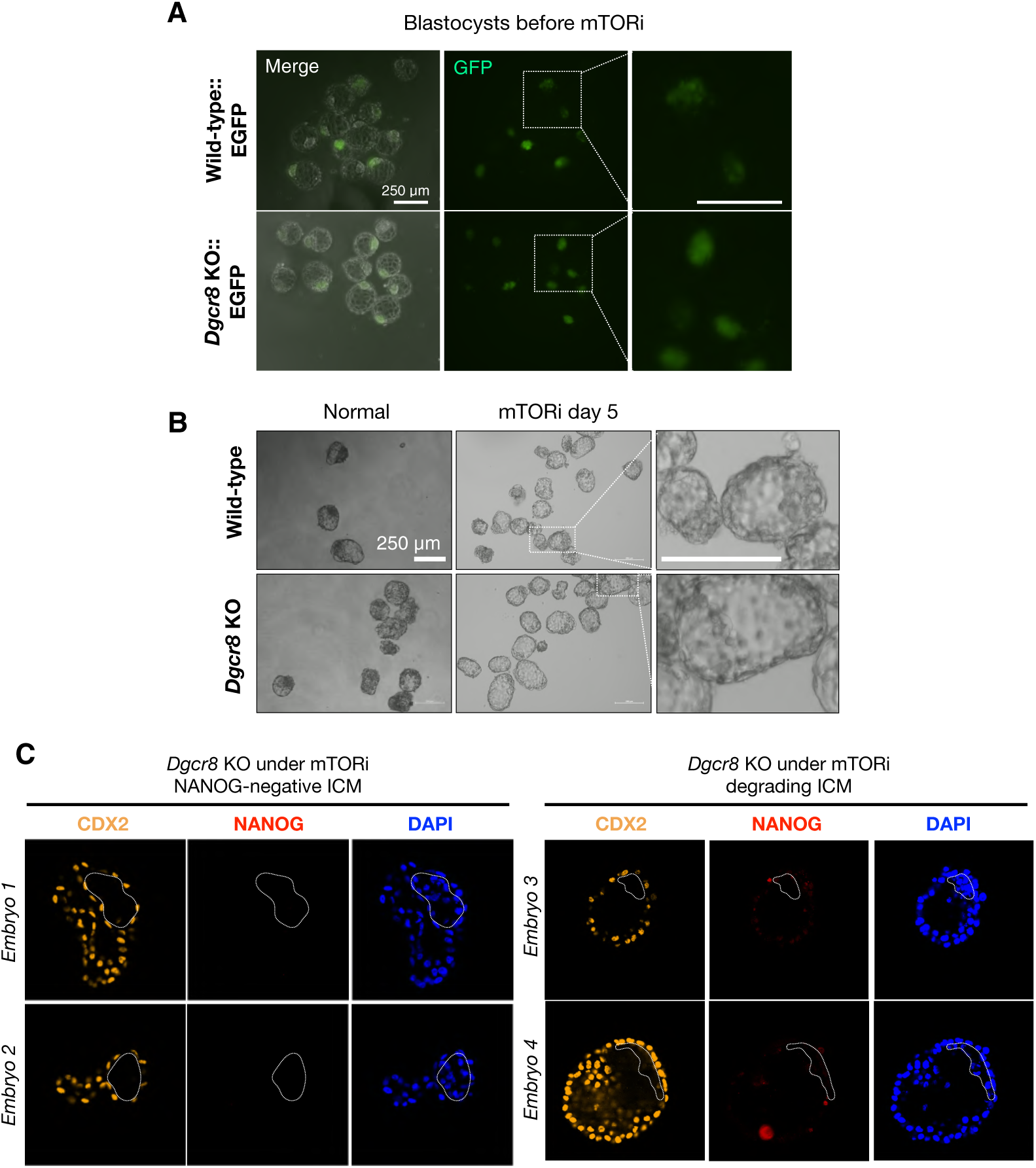
Characterization of chimeric wild-type and *Dgcr8* KO embryos. **A.** Bright field and/or fluorescence images of wild-type and *Dgcr8* KO chimeric embryos. EGFP signals shows the ESC contribution to the embryos. Both wild-type and *Dgcr8* KO ESCs contributed to the ICM in chimeric embryos, however, *Dgcr8* KO ESCs contributed more homogeneously. B. Bright field images of wild-type and *Dgcr8* KO chimeric embryos on day 5 of mTORi treatment. Right panels show higher magnification images of the area marked with white boxes. A representative *Dgcr8* KO embryo lacking a visibly large ICM is shown. C. Single z-stack images of *Dgcr8* KO blastocysts under mTORi. CDX2, NANOG, and DAPI stainings are shown. The dashed line marks the ICM or its remnants. The embryos retain blastocoel cavities.

**Figure S2.**
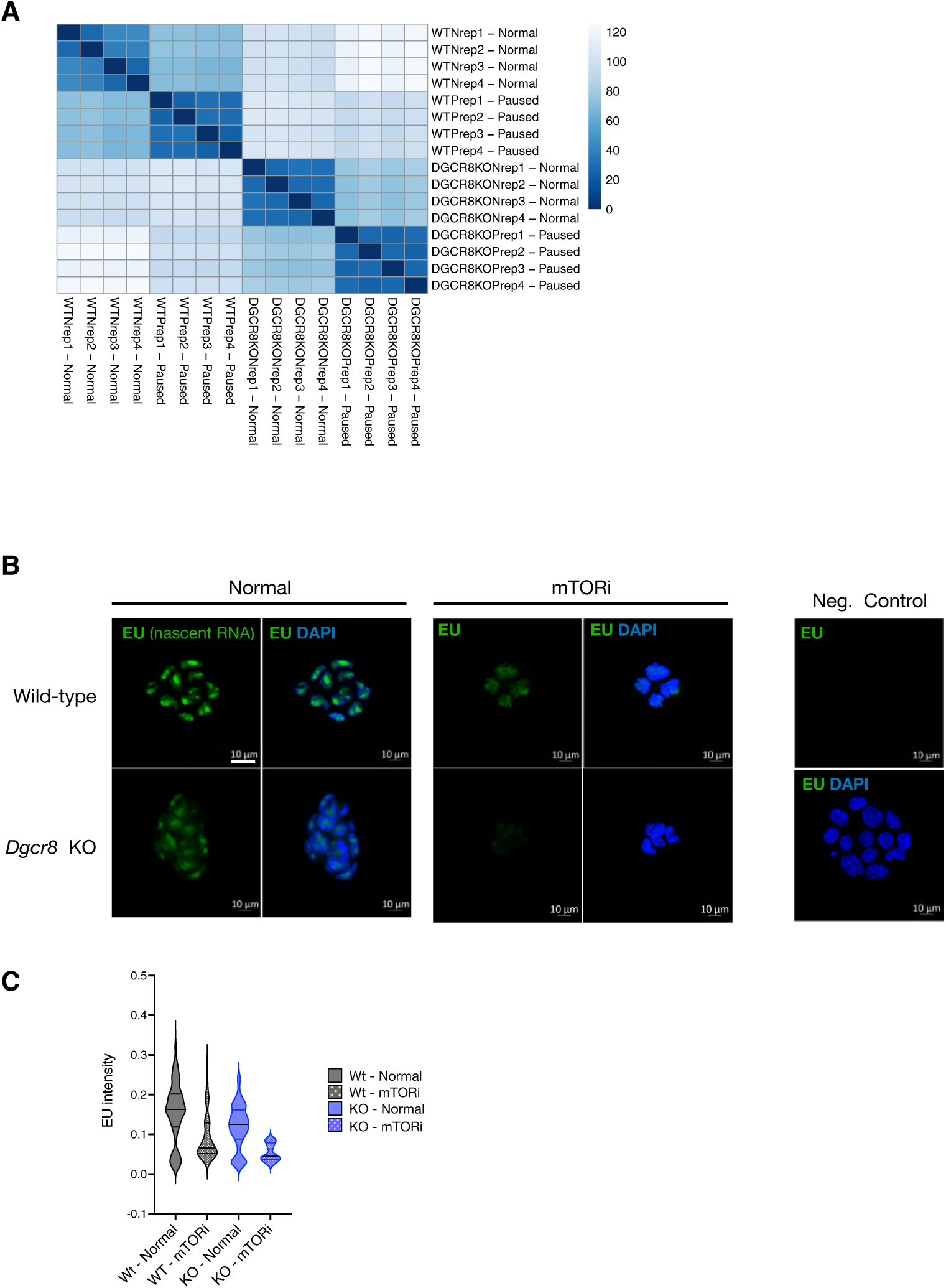
Further characterization of wild-type and *Dgcr8* KO ESC transcriptomes. **A.** Correlation heatmap of bulk RNA-seq datasets of wild-type and *Dgcr8* KO ESC under normal and mTORi conditions. **B.** Analysis of nascent RNA expression levels of wild-type and Dgcr8 KO ESCs in normal or pausing conditions. Cells were labeled with the nucleotide analog EU while live and visualized after fixing through conjugation of a fluorophore to EU via click chemistry. Negative control sample did not receive EU and was subject to the same procedure during all downstream steps. Scale bar = 20 um. **C.** Mean fluorescence intensity at single-cell resolution. CellProfiler was used to mark nuclei using DAPI as reference, and EU intensity was measured within the marker nuclei.

**Figure S3.**
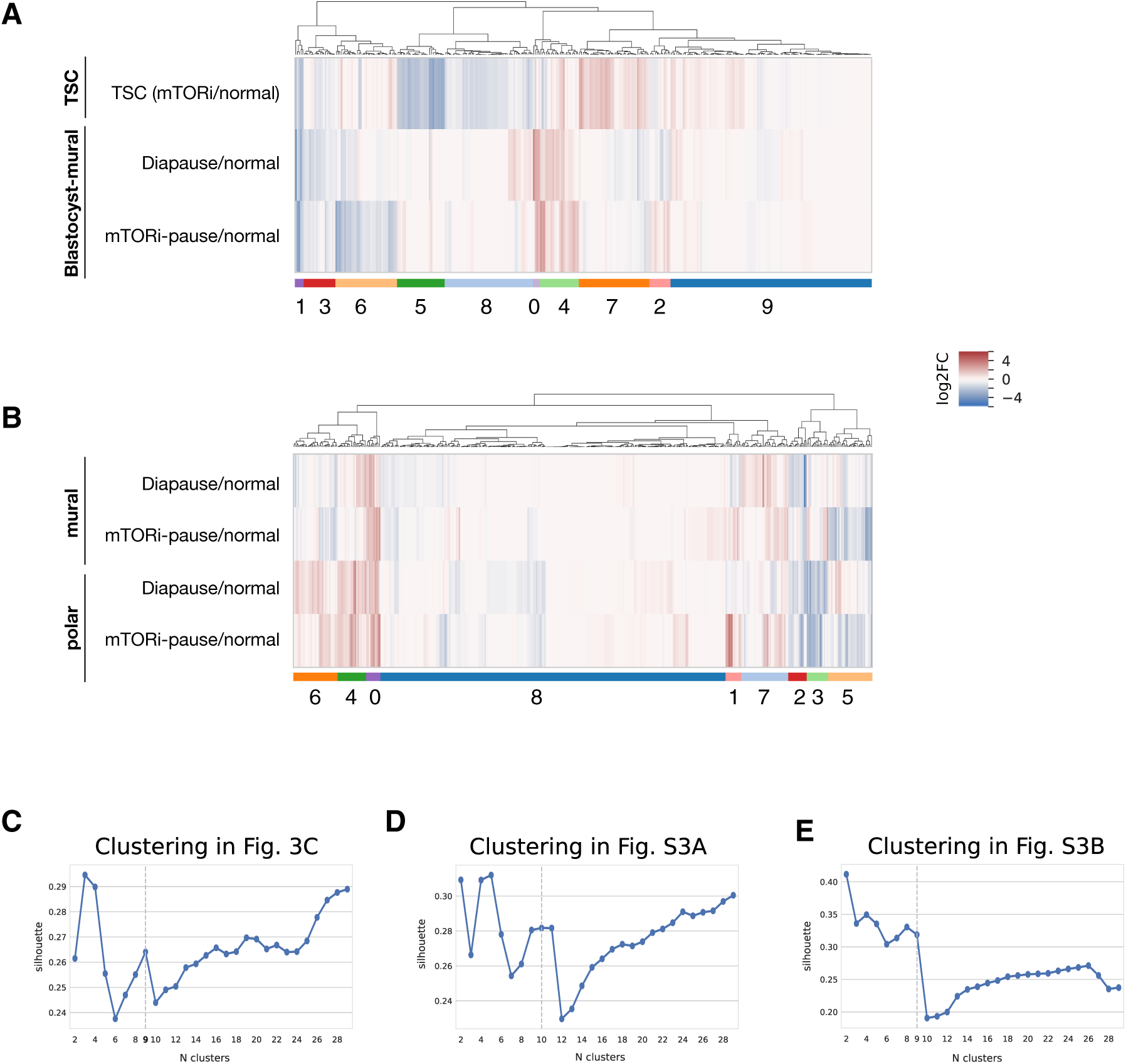
Details of miRNA clustering analysis. **A.** Heatmap showing miRNA expression changes in the mural side of diapaused embryos (in vitro and in vivo) compared to normal blastocysts, and in paused TSCs compared to normal TSCs. miRNAs were clustered into 10 clusters based on their expression levels in the three samples (see corresponding silhouette in Fig. S3C). Cluster 4 contains concordantly upregulated miRNAs (N=22), whereas clusters 3 and 1 contain concordantly downregulated miRNAs (N=18 and 5, respectively). **B.** Heatmap showing miRNA expression changes in the mural and polar sides of diapaused embryo compared to normal blastocysts. Cluster number 2 contains 10 concordantly upregulated miRNAs while cluster 2 contains 13 down-regulated miRNAs. **C.** Silhouette scores (measuring homogeneity within clusters compared to distance between clusters) for a range of cluster numbers from hierarchical clustering of log2FoldChange values from mTOR-inhibited ESC, mTOR-inhibited Blastocyst-polar cells, and diapaused Blastocyst-polar cells. The final number of clusters was identified as the best compromise of high-silhouette score and reasonable number of clusters. **D.** Silhouette of scores for a range of cluster numbers from hierarchical clustering of log2FC values for TSCs (mTORi/normal) and embryos (in vivo-and in vitro-diapaused blastocyst mural part/normal E3.5). **E.** Silhouette of scores for a range of cluster numbers from hierarchical clustering of log2FC values for diapaused embryos compared to normal blastocysts.

**Figure S4.**
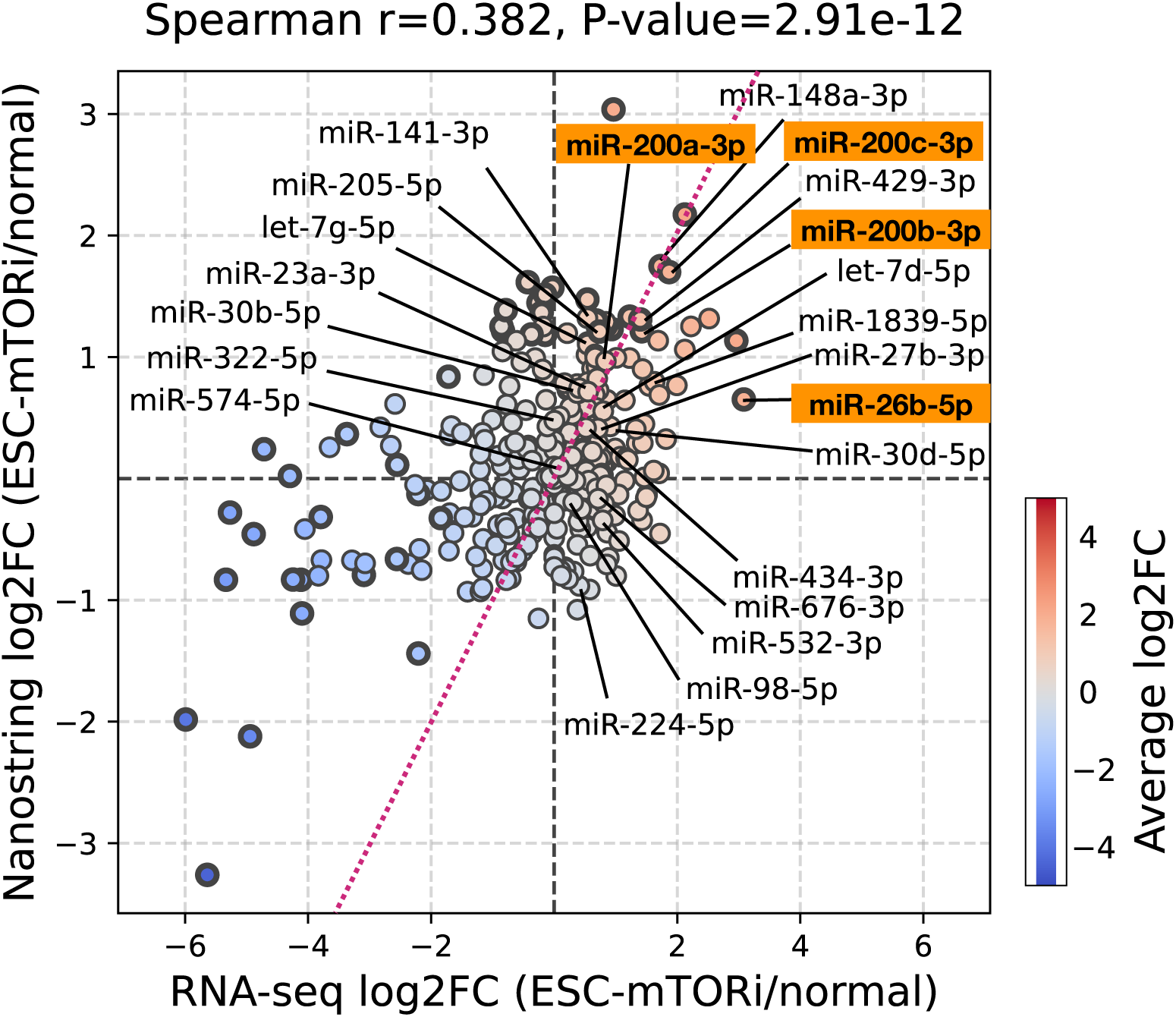
Amplification-free Nanostring analysis of miRNA expression. Comparison of miRNA expression levels resulting from small RNA-seq and Nanostring analyses. Overall, miRNA expression levels correlate, with miR-200 family showing the highest fold changes in both assays.

**Figure S5.**
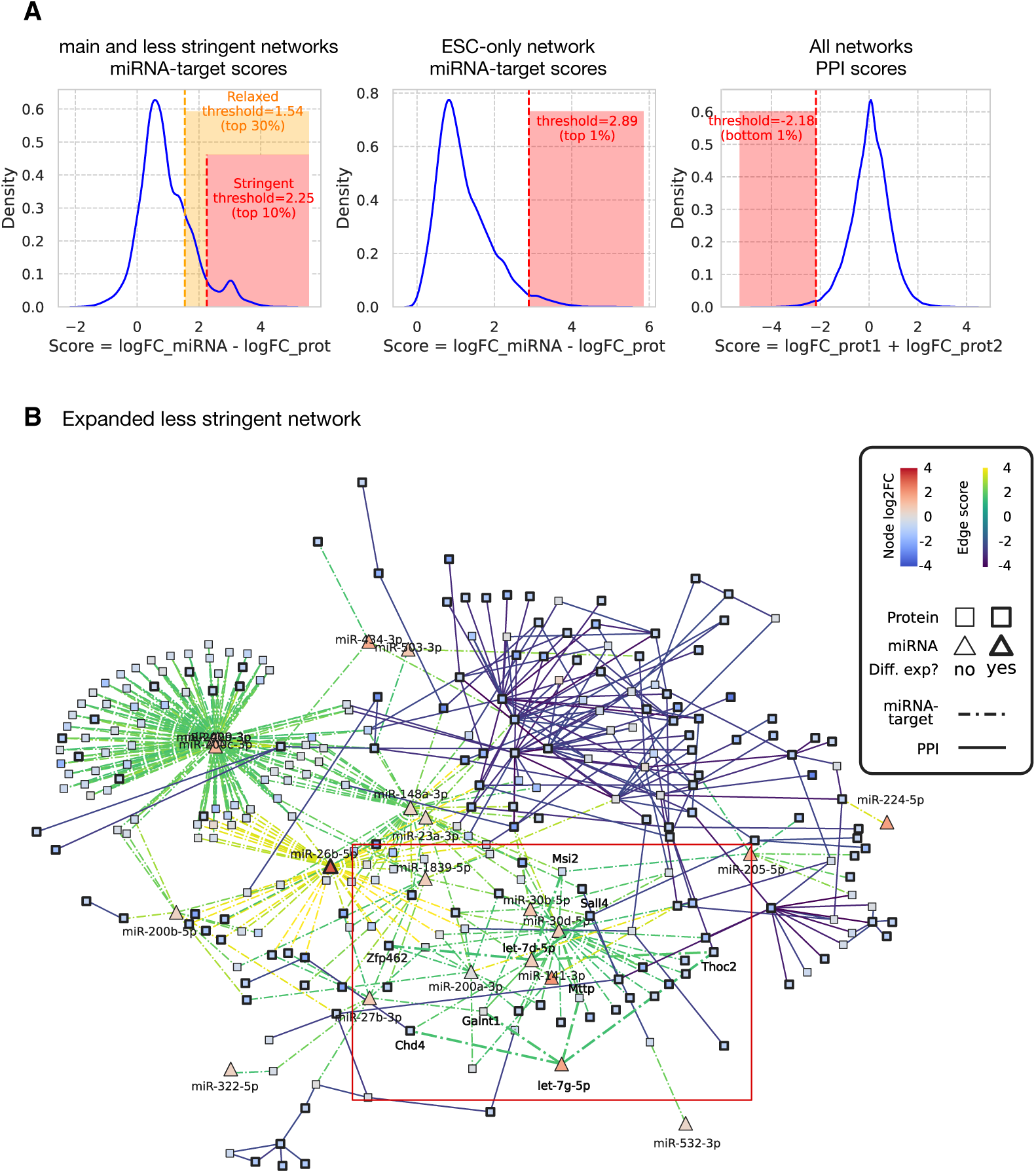
A less stringent network includes the let-7 miRNA. **A.** Network stringency thresholds. miRNA-target edges were scored as the difference in logFCs between the miRNAs and the target protein, while protein interactions were scored with the sum of logFCs. (Left) A subset of miRNAs was identified from their concordant up-regulated pattern across Blastocyst-polar pausing and ESC pausing, leading to a selection of miRNA-target edges with the visualized distribution of miRNA-target edge scores, from which two high-scoring sets were defined. (Middle) Similarly, a second subset of miRNAs was defined, from their positive logFCs values in mTORi ESC only, and top scoring edges were selected for. (Right) across all networks, protein-protein interactions were filtered for the bottom 1% lowest-scoring, indicative of co-down-regulation. **B.** An extended, less stringent miRNA-target network of dormancy. Let-7, a previously reported miRNA associated with mouse diapause^16^, is visible in this network, however does not pass the stringency threshold in the main network in Fig. 4C. A total of 8 target proteins are found, including significantly downregulated proteins Cdh4 (logFC=-1.107, adjusted-P-value=0.0048) and Thoc2 (logFC=-1.048, adjusted-P-value=0.017) targeted by both, while Sall4 (logFC=-0.862, adjusted P-value=0.0172) and Zfp462 (logFC=-0.791, adjusted-P-value=0.011) are additionally targeted by let-7-d.

**Figure S6.**
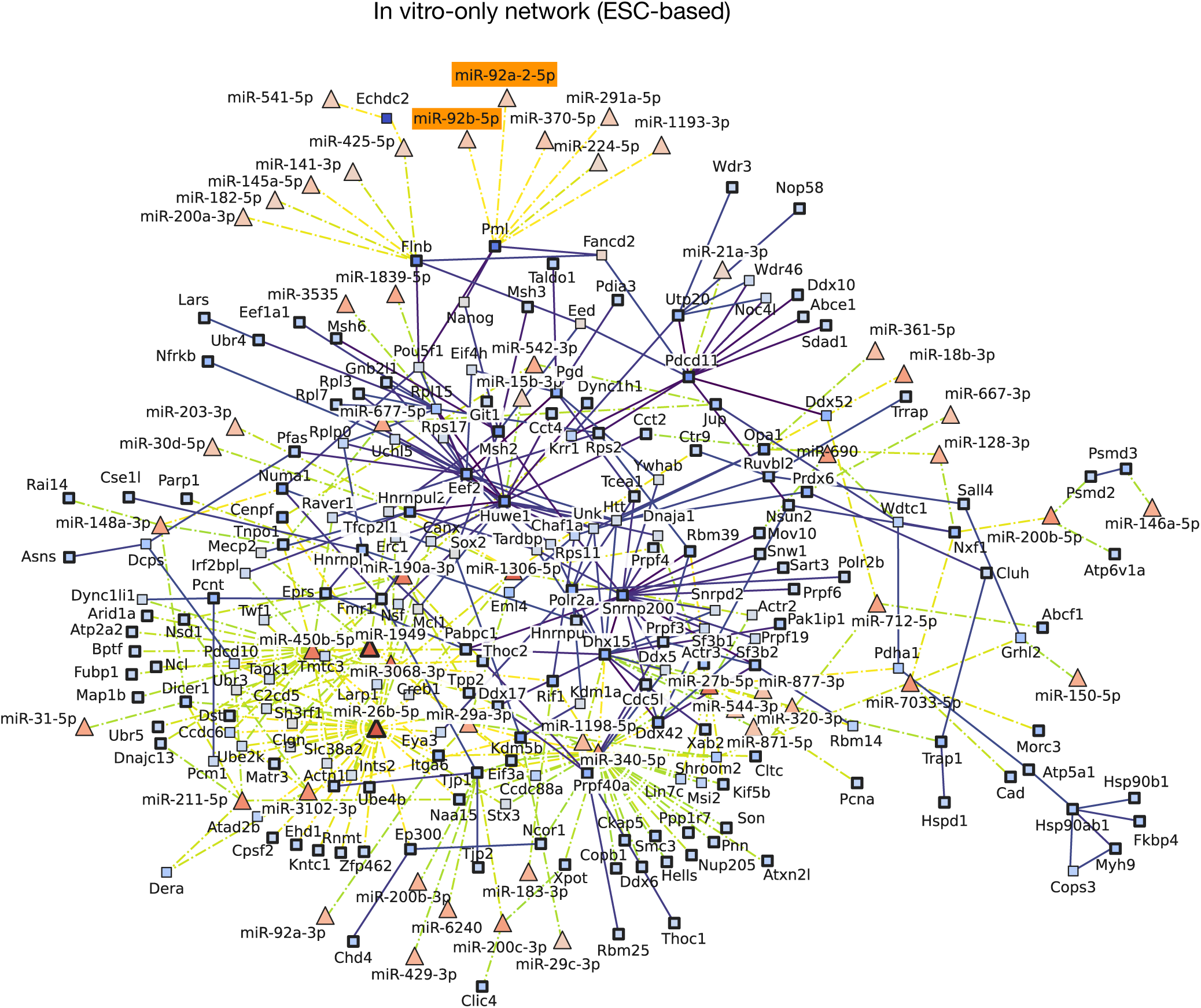
The miRNA-target network of ESC pausing. miRNA expression and proteome profiles of mTORi vs normal ESCs were used to construct the ESC-only network. The same stringency cutoff (top 10% of miRNA-target edges, and top 1% of PPIs) as the main network (Figure 4C) was applied. In addition to miR-200 family and miR-26, this network also recovers an evolutionarily conserved regulator of quiescence, miR-92^31^.

**Figure S7.**
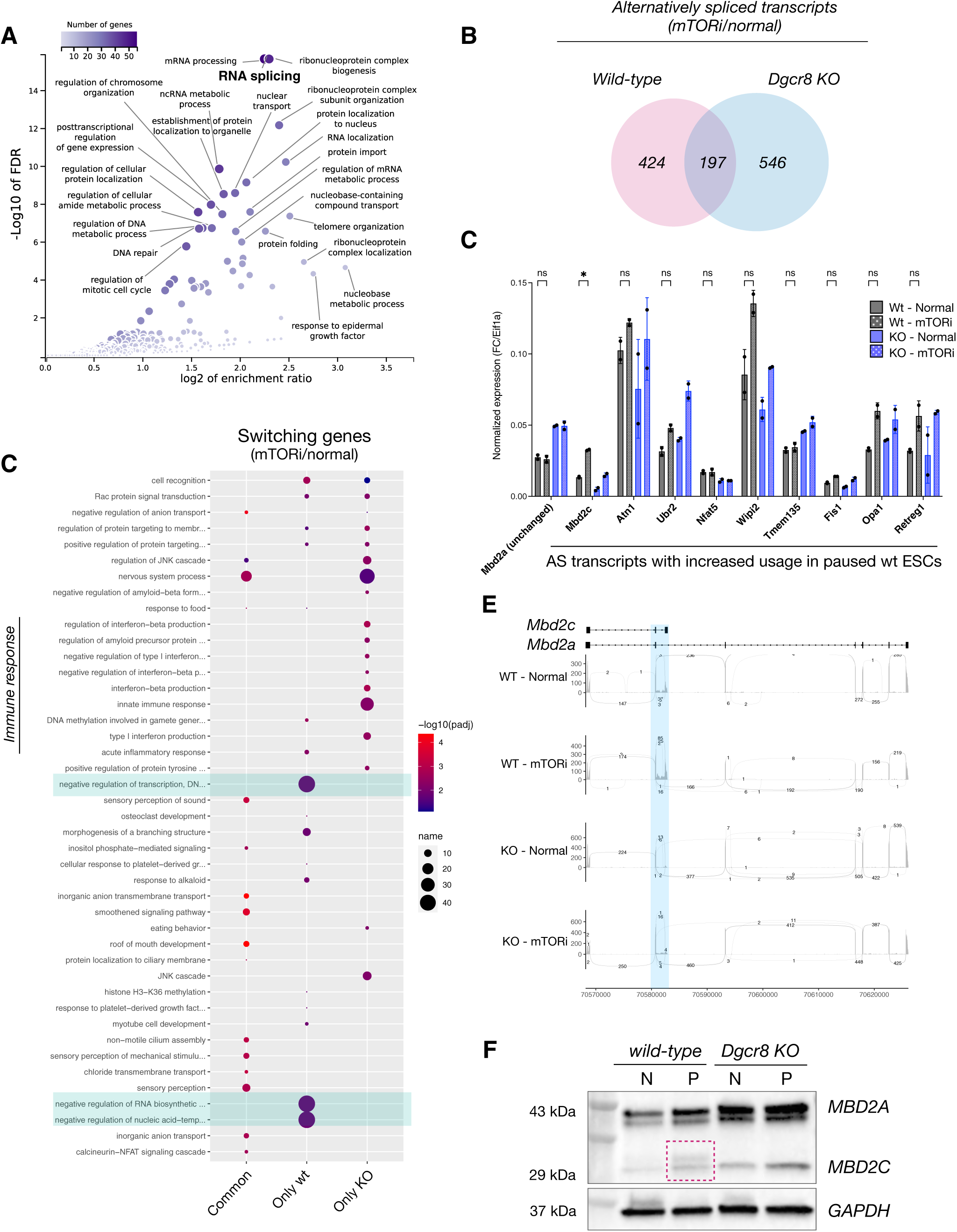
The transition into dormancy is associated with alternative splicing events. **A.** Gene ontology analysis of in-network proteins under putative miRNA control. RNA processing pathways comprise the most significantly altered pathways during diapause entry. RNA splicing is the top altered pathway. **B.** Numbers and overlap of alternatively spliced (AS) genes in wild-type and *Dgcr8* KO cells under normal and pausing conditions. **C.** Gene ontology analysis of alternatively spliced genes. Common and genetic background-specific pathways are shown. AS isoforms of negative regulators of transcription and RNA metabolism are enriched only in wild-type cells, whereas *Dgcr8* KO cells show an inflammatory response at this level. **D.** Selected AS isoforms and their validation by isoform-specific qPCR. Isoforms with increased usage in wild-type paused cells (compared to normal) were chosen. Primers were designed to amplify these isoforms specifically and are listed in the Methods section. **E.** Sashimi plot of the AS gene Mbd2. Arcs display the number of reads across splice junctions. **F.** Western blot against the MBD2 protein validates increased usage of the short isoform, Mbd2c in *Dgcr8* KO cells. An additional band around 32 kDa that may correspond to a posttranslational modification is visible in paused wild-type, but not *Dgcr8* KO ESCs. The MBD2 protein is expressed at higher than normal levels in *Dgcr8* KO cells.

**Figure S8.**
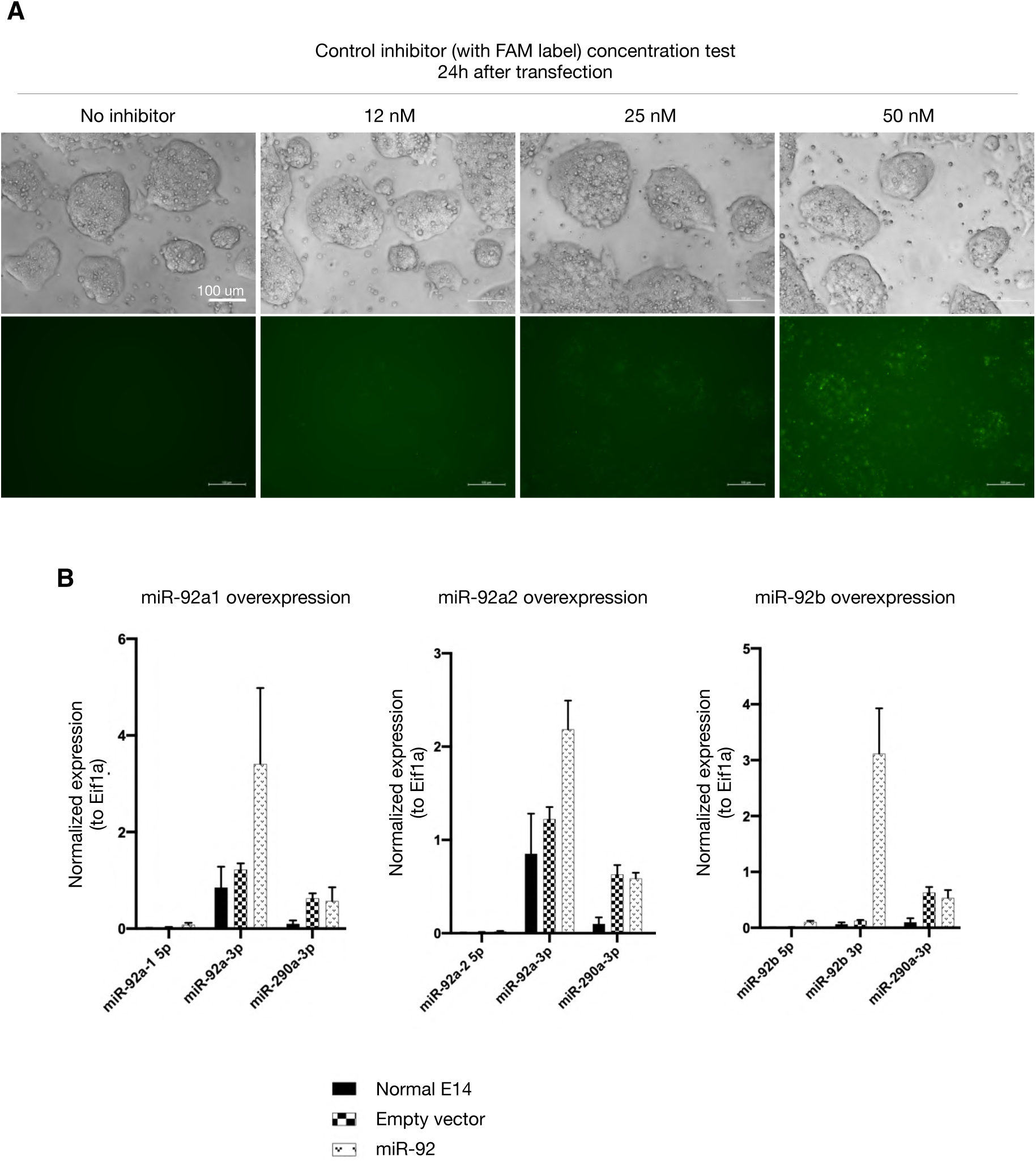
Characterization of miRNA inhibition and overexpression. **A.** Dilution series to determine the appropriate concentration for miRNA inhibitor treatments. FAM-labeled control inhibitor was used to visualize cellular intake. ESCs were transfected with the miRNA and imaged 24h later. 50 mM was chosen as the optimum concentration for use on embryos. Scale bar, 100 um. **B.** qPCR validation of miR-92 overexpression. Wild-type ESCs were transduced with virus carrying miR-92a1/a2/b or the corresponding empty vector. Expression levels of processed miRNAs (miR-92a-3p or miR-92b-3p) were assessed using specific TaqMan probes. miR290a-3p was used as control. Of note, miR-92a1 and miR-92a2 3p arms have the same sequence.

**Figure S9.**
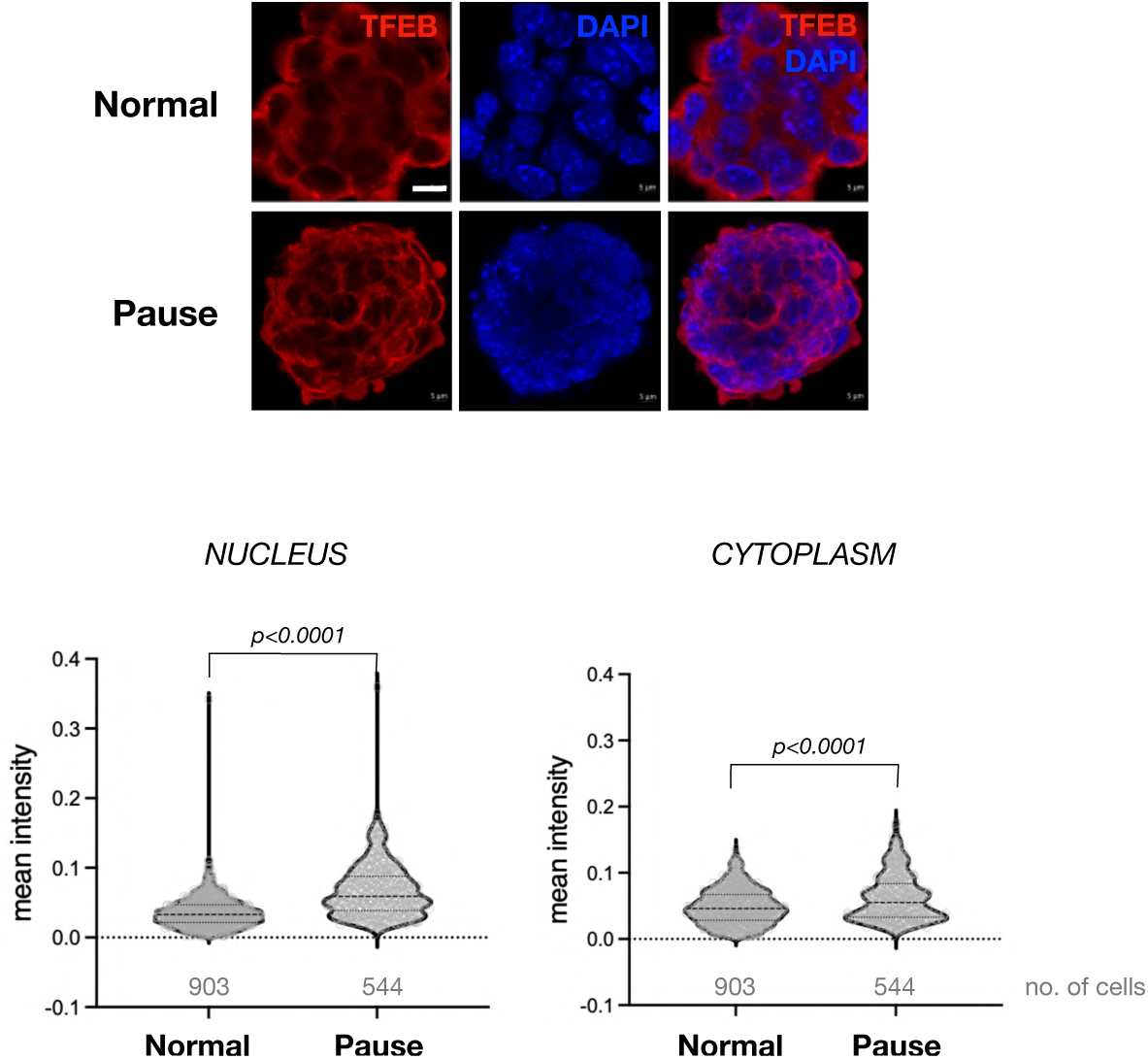
TFEB expression and subcellular localization. TFEB staining in wild-type normal and paused ESCs. Lower panels show single-cell quantifications of mean fluorescence intensity in the nucleus and cytoplasm. Scale bar = 5 µm. Statistical test is Kolmogorov-Smirnov test, two-sided.

## METHODS

### Animal experimentation

All animal experiments were performed according to local animal welfare laws and approved by local authorities (covered by LaGeSo licenses ZH120, G0284/18, and G021/19). Mice were housed in ventilated cages and fed ad libitum.

### Cell lines and culture conditions

#### Mouse ESCs

Wild-type and *Dgcr8* KO^24^ E14 ESCs (received from S. Kinkley Lab, MPIMG and Constance Ciaudo Lab, ETH respectively) were used. Cells were plated on 0.1% gelatin-coated dishes and grown in DMEM high glucose with Glutamax media (Thermo Fisher Scientific, 31966047) supplemented with 15% FBS (Thermo Fisher Scientific, 2206648RP), 1x NEAA (Thermo, 11140-035), 1x b-mercaptoethanol (Thermo Fisher Scientific, 21985023), 1x Penicillin/streptomycin (Life Technologies, 15140148) and 1000 U/mL LIF and grown at 37°C in 20% O2 and 5% CO2 incubator. At each passage, cells were dissociated using TrypLE (Thermo Fisher Scientific, 12604-021) with media change every day. Cells tested negative for mycoplasma.

#### Mouse TSCs

TSCs (received from M. Zernicka-Goetz Lab) were grown on mitotically-inactivated mouse embryonic fibroblasts in media containing RPMI1640 + GlutaMAX (Thermo Fisher Scientific, 61870010), 20% FBS, b-mercaptoethanol (Thermo Fisher Scientific, 21985023), 1x Penicillin/ streptomycin (Life Technologies, 15140148), 1x sodium pyruvate, 25 ng/µl FGF4 (R&D Systems, 235-F4-025) and 1 µg/ml Heparin (Sigma, H3149). Cells were depleted off feeders before collecting for analysis. Cells tested negative for mycoplasma.

### Developmental pausing setup

#### Mouse ESC/TSC pausing

ESCs and TSCs were treated with the mTOR inhibitor INK-128 at 200 nM final concentration. Media was replenished as required. ESCs were paused for 6 days and TSCs for 12 days before collection of cells for small RNA profiling.

#### In vitro diapause

Blastocyst-stage embryos were cultured in KSOM+AA medium (Sigma, MR-101) and treated with 200 nM of RapaLink-1 to induce diapause *in vitro*.

#### In vivo diapause

In vivo diapause was induced as previously described^6^ after natural mating of CD1 mice. Briefly, pregnant females were ovariectomized on E2.5 and afterwards injected with 3 mg medroxyprogesterone 17-acetate subcutaneously (Sigma, M-1629) on E3.5 and E5.5. Diapaused blastocysts were flushed from uteri in M2 media after 4 days of diapause at EDG7.5.

### Morula aggregations

Wild-type diploid morulae collected from CD1 (Hsd:ICR) females were used for aggregations. Wild-type-EGFP or *Dgcr8* KO-EGFP E14 mouse ESCs were combined with morulas as described^54^. Cells express an EGFP gene expressed from a CAGGS promoter and randomly integrated into the genome as a marker of ESC origin.

### Bulk RNA-seq

800,000 wild-type or *Dgcr8* KO^24^ mouse ESCs in normal or mTORi culture (day 2) were used. Total RNA was extracted using the RNeasy extraction kit (Qiagen, 74104). 1 µg of total RNA was used for standard library preparation using NEBNext PolyA mRNA magnetic isolation module (NEB, E7490). In brief, polyA mRNA isolation was done followed by first-strand and second-strand cDNA synthesis. Double-stranded cDNA was purified using AMPure beads (Beckman Coulter, A63881). Adaptor ligation was performed in a 5-fold diluted ice-cold adaptor dilution buffer. PCR enrichment of adaptor-ligated DNA was done using the following conditions: Initial denaturation - 98C for 30 seconds; Denaturation - 98C for 10 seconds; Annealing - 65C for 75 seconds (8 cycles); final extension - 65C for 5 minutes. Purification of the PCR reaction was done using AmPure beads. Library quality was assessed by running samples on Tape Station (Agilent, 4150).

### Small RNA-seq on ESCs and TSCs

#### Sample preparation

Small RNA was extracted using miRNeasy kit (Qiagen, 217004). In brief, the cell pellet was homogenized in Qiazol and incubated on a benchtop for 5 min. Chloroform was added to the homogenate and was shaken vigorously for 15 sec and incubated at room temperature for 3 min. The homogenate was centrifuged at 12000 g for 15 mins at 4C. To the aqueous layer, 70% ethanol was added and shaken vigorously. The sample was then pipetted into the RNeasy Mini spin column and centrifuged at room temperature. The flow-through contains the miRNA and other small RNA. To the flow-through, 100% ethanol was added and shaken vigorously. The sample was pipetted into an RNeasy MinElute spin column (Qiagen Cat No: 74204). The column was washed with buffers RWT, RPE, and 80% ethanol. Small RNAs were extracted in 14ul of RNase-free water.

#### Library preparation and sequencing

10 ng RNA was used as input. Small RNA libraries were prepared using the SMARTer smRNA-seq Kit for Illumina (Takara, 635031) with 12 amplification cycles followed by bead size selection (ratio 0.95x-0.7x), followed by an additional clean-up step with the bead-to-sample ratio of 0.95x. Library QC was done using Qubit and BioAnalyzer and the final libraries were quantified using KAPA Library Quantification Kit (Roche, 07960140001). Libraries were diluted to 2 nM final concentration for pooling. Pooled libraries were sequenced on a HiSeq4000 flowcell with single-end 75 bp reads to get ∼15M reads per sample.

### Ultra low-input small RNA-seq on dissected embryos

#### Sample preparation

Normal blastocysts at E4.5, in vivo paused blastocysts at EDG7.5, and mTORi-treated blastocysts were used for microdissection. Laser-assisted separation of ICM and mural trophoblast was performed using a Hamilton Thorne XYRCOS laser system with power set to 100% and a pulse length of 300 µs (see Supplementary Video 1). For dissection, blastocysts were fixed between two holding pipettes. While the holding pipettes were carefully pulled apart, cellular connections were cut by successive laser pulses, allowing the separation of the blastocyst into pieces containing ICM or mural trophectoderm. Immediately after separation the tissue fragments were transferred into 7 µl PBS and frozen down on dry ice.

#### Library preparation and sequencing

Small RNA libraries were prepared using the “TrueQuant small RNA-Seq” kit from GenXPro (Germany) according to the manual. The method is based on a single-tube protocol for ligation of UMI-containing adapters to RNA molecules followed by Reverse Transcription and PCR^55^. No size selection of the small RNAs was performed. Small RNAs were amplified with 12-20 cycles to reach the required amounts for sequencing. Sequencing was performed on an Illumina NextSeq500 instrument with 1x75 bps.

### Small RNA-seq data analysis

#### Mapping and quantification

The miRDeep2^57^ (v2.0.1.2) algorithm was applied for quantification of miRNAs across samples, using the reference annotations of mature and hairpin miRNAs from miRBase^58^ (version 22). The steps of the algorithm include preprocessing and quality control of reads, alignment of reads (mapper.pl script, options -j -l 17, other options set to default) with ambiguous reads assignment and multi-loci reads processing (default: maximum of 5 multi-site mapping), and quantification of miRNAs against miRBase mouse mature and hairpin sequences, after filtering for other non-coding RNAs. Across samples, an average of 44.5% of reads were aligned. The quality of quantification was confirmed by evaluating Pearson correlation of read counts between replicates for each set of conditions across experiments, observing an average correlation of 0.944.

#### Differential expression analysis

Differential expression analysis was applied for each experiment using DESeq2^59^, comparing each set of condition samples against the control samples for both samples representing embryonic fate and samples representing extraembryonic fate separately. Low read count miRNAs were discarded by keeping only those miRNAs showing a normalized read count above 5 in at least 3 samples. Read-counts quantification was transformed into regularized log counts for downstream analyses such as PCA visualization of samples. Differentially expressed genes were identified with thresholds |logFoldChange|≥log2(1.5) (representing an increase or a decrease of at least half the value in the control samples) and adjusted p-value<0.1 (Benjamin-Hochberg correction).

#### Differential expression status agreement and clustering

From the results of DESeq2 differential expression analyses across experiments, logFoldChange values of miRNAs from each analysis were aggregated into a single table. The concordance of differential expression status for each miRNA was evaluated for each group of experiments by comparing the logFoldChange sign and magnitude. To systematically identify groups of miRNAs with concordant patterns across ESCs and embryo experiments, we applied hierarchical clustering to the miRNA fold change profiles (with Ward’s linkage for clusters distance and Euclidean distance between pairs of miRNAs). We calculated the average silhouette coefficient, which measures the homogeneity within clusters versus the distance between clusters) for an increasing number of clusters (2 to 30), and selected the final number by maximizing this silhouette score, balanced by the number of clusters. The final indexing of clusters per experiment reflects the number of miRNAs within the cluster, from 0 the smallest cluster to N the largest cluster. Clustering was performed for the following groups of experiments separately: ESCs versus Blastocyst-polar, TSCs versus Blastocyst-mural, and Blastocyst-polar versus Blastocyst-mural. For each group, average logFoldChange values were calculated for each cluster, and clusters grouping miRNAs with concordant logFoldChange (either positive or negative across all comparisons) were extracted for downstream regulatory network analyses.

### Nanostring

#### Quantification

Total RNA was isolated using the Rneasy extraction kit (Qiagen, 74104). 150 ng of total RNA in a volume of 3 µl was used as the starting material. miRNA sample preparation and miRNA CodeSet hybridization were done following the manufacturer’s protocol (MAN-C0009-07). The Sprint cartridge was then run on nCounter® Sprint Profiler. Quantification data was generated using nSolver from Nanostring. Of the four housekeeping genes, B2m was barely expressed across all samples, so normalization was done only using the following 3 housekeeping genes : Actb, GAPDH, and Rpl19. A total of four samples were quantified: ESC-WT (2 replicates, samples 01 and 02) and mTORi paused ESC (2 replicates, samples 05 and 06).

#### Differential expression

RCC values were extracted so as to be used for differential expression using DESeq2 with RUV correction^60^. This correction method allows for exploring and removing components of unwanted variations detected in the data. The first component of unwanted variation was removed. From the available housekeeping genes, B2m was confirmed to be too little expressed to be used for housekeeping genes normalization. Actb appeared as differentially expressed in ESCs on day 6 of pausing, but was still included as housekeeping for normalization. DESeq2 is then applied to the transformed values of the measured mouse miRNAs. Differentially expressed miRNAs were identified using the following thresholds: absolute log2FC greater or equal to log2(1.5), and adjusted P-value lower or equal to 0.1.

### Global proteomics

#### Sample preparation

Proteomics sample preparation was done according to a published protocol with minor modifications^61^. In brief, 5 million cells in biological duplicates were lysed under denaturing conditions in 500 µl of a buffer containing 3 M guanidinium chloride (GdmCl), 10 mM tris(2-carboxyethyl)phosphine, 40 mM chloroacetamide, and 100 mM Tris-HCl pH 8.5. Lysates were denatured at 95°C for 10 min shaking at 1000 rpm in a thermal shaker and sonicated in a water bath for 10 min. 100 µl lysate was diluted with a dilution buffer containing 10% acetonitrile and 25 mM Tris-HCl, pH 8.0, to reach a 1 M GdmCl concentration. Then, proteins were digested with LysC (Roche, Basel, Switzerland; enzyme to protein ratio 1:50, MS-grade) shaking at 700 rpm at 37°C for 2h. The digestion mixture was diluted again with the same dilution buffer to reach 0.5 M GdmCl, followed by a tryptic digestion (Roche, enzyme to protein ratio 1:50, MS-grade) and incubation at 37°C overnight in a thermal shaker at 700 rpm. Peptide desalting was performed according to the manufacturer’s instructions (Pierce C18 Tips, Thermo Fisher Scientific). Desalted peptides were reconstituted in 0.1% formic acid in water and further separated into four fractions by strong cation exchange chromatography (SCX, 3M Purification, Meriden, CT). Eluates were first dried in a SpeedVac, then dissolved in 5% acetonitrile and 2% formic acid in water, briefly vortexed, and sonicated in a water bath for 30 seconds prior to injection to nano-LC-MS/MS.

#### Run parameters

LC-MS/MS was carried out by nanoflow reverse phase liquid chromatography (Dionex Ultimate 3000, Thermo Fisher Scientific) coupled online to a Q-Exactive HF Orbitrap mass spectrometer (Thermo Fisher Scientific), as reported previously62. Briefly, the LC separation was performed using a PicoFrit analytical column (75 µm ID × 50 cm long, 15 µm Tip ID; New Objectives, Woburn, MA) in-house packed with 3-µm C18 resin (Reprosil-AQ Pur, Dr. Maisch, Ammerbuch, Germany).

#### Peptide analysis

Raw MS data were processed with MaxQuant software (v1.6.10.43) and searched against the mouse proteome database UniProtKB with 55,153 entries, released in August 2019. The MaxQuant processed output files can be found in Table S6, showing peptide and protein identification, accession numbers, % sequence coverage of the protein, q-values, and label-free quantification (LFQ) intensities.

### Differential expression analysis and enrichment analyses

The protein levels across the three replicates of each condition were extracted from the reported label-Free Quantification values from the mass spectrometry experiment. Protein intensities were scaled by the median value per sample. No proteins were removed for the differential expression analysis. The analysis was performed using Limma63 and Limma-Voom^64^. We applied a threshold of | logFoldChange|≥log2(1.5) and a threshold of adjusted P-value>0.1 for the identification of differentially expressed proteins. GO-term and pathway enrichment analyses were performed with the online tool WebGestalt^65^, using the non-redundant sets of GO terms, and the KEGG pathways. Default parameters were kept, notably for filtering categories (minimum number of genes: 5 per category, maximum number: 2000) and for correction for multiple testing (Benjamini-Hochberg correction, FDR<0.05 for filtering significant categories).

### miRNA-target network

Predictions of miRNA-targets were gathered from the miRDB database (V6.0), annotated into three categories of confidence (low, medium, high) following the official documentation on the confidence score (associated thresholds: ≥50, ≥60, ≥80). Only medium and high-confidence miRNA-target interactions were retained for downstream analysis. In addition, experimentally validated interactions from miRTarBase V8.0 were included, retaining only functional interactions. This led to a total of 705,242 edges (664,660 from miRDB, 30,795 from miRTarBase, 9,787 from both) between 1,998 miRNAs and 17,802 proteins. Annotation of the edges and their nodes was done using the log2FoldChange values obtained from miRNAs and proteomics differential expression analyses comparing mTORi paused ESC against controls. We established a score for the edges defined as the difference of log2FoldChange values between the miRNA and the target measured protein. Such score emphasizes the expected negative regulation of miRNAs onto their target translation so that a highly up-regulated miRNA linked to a highly down-regulated protein will lead to a highly positive edge score. This resulted in a subset of 80,099 edges with annotated scores, for 558 miRNAs and 5,019 targets. From these edges, three subnetworks were built after filtering for connections between miRNAs and proteins with non-zero logFoldChange values. First, subsets of miRNAs of interest were defined based on their differential expression values either from the mTOR-inhibited paused ESC analysis, or from their joint patterns evaluated across pausing experiments of ESC and Blastocyst-polar cells. The first network consists of regulatory relationships between the 24 concordantly upregulated miRNAs between ESC and embryos and their targets., yielding 5,234 miRNA-target edges with 2,104 unique proteins. As the 1% of top-scoring edges yielded only 3 miRNAs, we relaxed the threshold down to the top 10%, yielding a subnetwork of N=196 edges between 17 unique miRNAs and 172 proteins. The second network consisted of the same 24 miRNAs, relaxing the score threshold down to the top 30% of miRNA-target regulatory interactions. This allowed us to explore additional regulatory relationships that do not exhibit strong regulatory patterns (i.e. strong target downregulation upon miRNA upregulation), and so discover miRNA-target subnetworks with milder phenotypic changes, but nonetheless putatively still important in the context of pausing. We finally constructed a third network from the ESC experimental data only, considering the full set of miRNAs with a positive logFoldChange from the differential expression analysis of mTOR-inhibited ESCs. As this resulted in a large network, we selected and inspected for downstream analysis only the top 1% of highest-scoring miRNA-target connections, yielding a subnetwork of N=355 edges with 55 unique miRNAs and 266 proteins.

### Multipartite miRNA-protein and protein-protein networks

To address how perturbation of miRNAs propagates downstream and which other proteins or subnetworks are impacted by the deregulated miRNAs, we extended the miRNA-target network to a multipartite network of miRNA-targets and protein-protein interactions assembled from multiple sources of gene connections. The use of multiple sources aimed at both covering more relationships than any single resource, as well as identifying connections with multiple supporting sources. Protein-protein interactions (PPIs) were gathered from the StringDB database (V12.0), filtered so as to keep the top 10% of all interactions after sorting on the combined score, and dropping those reported from text-mining only. These were complemented with interactions reported in the BioGrid database (V4.4.224). This represented a total of 84,363 deduplicated, high-confidence interactions, involving 12,896 unique proteins. The nodes in this protein-protein network were annotated with the logFoldChange values obtained from the differential proteomic analysis upon mTOR-inhibited paused ESCs. As for the miRNA-target edges, protein-protein interactions were scored using the logFoldChange values of the nodes they connect, where we defined the edge score between two proteins as the sum of the logFC of two connected proteins, highlighting the expected co-expression of interacting proteins. This yielded a total of 34,846 protein-protein edges including 4,730 unique proteins with measured logFCs. From these, the top 1% highest-scoring edges were retained and included in each of the different miRNA-target subnetworks, resulting in a multipartite network of the most important regulatory interactions, supposedly representing the core of the dormancy network. Of note, to enhance the clarity of visualization of the resulting networks, we removed protein nodes where the protein had only one connection (miRNA or protein) while not significantly down-regulated.

### miRNA loss-of-function in blastocysts via miRNA inhibitor injections

Inhibitors (antimiRs) against miR26b-5p and miR200a/b/c-3p were purchased from Qiagen (339130 and 339160 respectively). AntimiRs in injection media (1mM Tris-HCL pH 7.5 and 0.5 mM EDTA in embryo grade water) were injected into each cell of 2-cell-stage embryos (BL/6xCast hybrid) using Leica DMIRB inverted microscope, Narishige MMO micromanipulators/microinjectors, Eppendorf Celltram 4r microinjector and Eppendorf FemtoJet microinjector. Embryos were then transferred to fresh KSOM+AA media (Sigma, MR-101) and cultured until the early blastocyst stage. Blastocysts were split into two groups and treated either with DMSO or 200 nM RapaLink-1. As negative control, a non-targeting inhibitor (Qiagen, 339136) with no sequence hits of >70% homology to any organism in the NCBI and miRBase databases was used. The survival rate and duration of embryos in mTORi-induced pausing were scored.

### miR-92 overexpression

Pre-miR sequences of miR-92a and miR-92b were PCR amplified and cloned into pLKO.1-puro lentiviral vector between AgeI and EcoRI restriction sites. Virus production and transduction were done using standard protocols. In brief, 4ug of lentiviral plasmids were packaged with equal parts of viral packaging vectors (pVSV-G, pMDL, pRSV) by transfecting them in HEK 293T cells. Virus production was carried out for 72 hours. The final precipitated virus was then suspended in 200 ul ice-cold PBS. 30ul of viral suspension was then used to infect 100,000 mESCs in single-cell suspension for 1 hour at 37C. After this, the infected mESCs were seeded onto a 6-well culture plate. The media was changed the next day. Puromycin selection was done 48h after infection and overexpression of respective microRNAs was assessed by miRNA qPCR using Taqman probes against miR-92a-5p and 3p; miR-92b-5p and 3p; miR-290a-3p (positive control). The relative microRNA expression was calculated against the eIF1alpha housekeeping gene.

### Immunofluorescence (IF)

#### ESCs

Cells were cultured on glass coverslips and were fixed in 4% PFA for 10 min at room temperature, washed once in PBS, and then permeabilized with 0.2% Triton-X100 in PBS for 5 min on ice. After washing once in PBS-T (PBS with 0.2% Tween-20), cells were blocked with blocking buffer (PBS-T, 2% BSA, and 5% goat serum (Jackson Immunoresearch/Dianova, 017-000-121) for 1 hour at room temperature. Cells were then stained with the primary antibodies PML, SC-35, DDX-6 overnight at 4C. The cells were washed thrice with wash buffer (PBS-T, 2% BSA) for 10 min. Anti-mouse 488 or anti-rabbit 568 secondary antibody conjugated with Alexa Fluor was added to cells at a dilution of 1:700 in blocking buffer and incubated for 1 hour at room temperature, followed by 3 washes with wash buffer for 10 min. The coverslips were then mounted with Vectashield with DAPI (Vector labs, H-2000) and sealed with nail polish. Imaging was done using a Zeiss LSM880 Airy microscope using Airy scan mode and image processing was done using Zen black and Zen Blue software (version 2.3). Image quantification was done using CellProfiler (version 4.2.1) where nuclei or cells that were denoted as primary objects were identified and the normalized intensities of the respective protein stained were measured against nuclear or cell area (https://cellprofiler.org/). Data were plotted using GraphPad Prism (version 9).

#### Embryos

Embryos were fixed for 10 minutes in 4% PFA, then permeabilized for 15 minutes in 0.2% Triton X-100 (Sigma, T8787) in PBS. Permeabilized embryos were incubated in blocking buffer (0.2% Triton X-100 in PBS + 2% BSA Fraction V 7.5% (Thermo Fisher Scientific, 15260-037) + 5% goat serum (Jackson Immunoresearch/Dianova, 017-000-121) for 1h. Then embryos were incubated overnight at 4C with the CDX-2 and NANOG primary antibodies. Primary antibodies were diluted in blocking buffer, Prior to incubation with the secondary antibodies, embryos were washed three times in washing buffer (0.2% Triton X-100 in PBS + 2% BSA). Then, the embryos were incubated with the secondary antibodies for 1h at room temperature. The donkey anti-mouse Alexa Fluor 680 and donkey anti-rabbit Alexa Fluor 568 antibodies were diluted 1:200 in blocking buffer. After incubation with the secondary antibody, embryos were washed three times in washing buffer. Then, embryos were mounted on a microscope slide with a Secure-Seal™ Spacer (8 wells, 9 mm diameter, 0.12 mm deep, Thermo Fisher Scientific, S24737), covered with a cover glass, and sealed with nail polish.

Imaging was done using a Zeiss LSM880 Airy microscope using Airy scan mode and image processing was done using Zen black and Zen Blue software (version 2.3). Image quantification was done using CellProfiler^65^ (version 4.2.1) where nuclei or cells that were denoted as primary objects and the normalized intensities of the respective protein stained were measured against nuclear or cell area (https://cellprofiler.org/). Data were plotted using GraphPad Prism (version 9).

### Transcription Factor Binding Site (TFBS) mining at candidate miRNA promoters

To identify potential transcription factors regulating the set of concordant up-regulated miRNAs, we performed a motif search over the promoter sequences of these miRNAs. Promoters were taken from the work of de Rie et al. 2017^34^, totalling 1,468 promoter regions (lifted over to mm10 genome assembly), each associated to one miRNA. Overlapping regions were merged, resulting in 641 sequences. Larger regions were then identified by extending from the 3’ end by 1,500 nucleotides upstream and 500 nucleotides downstream. Motifs of transcription factors were obtained from the JASPAR database^66^, taking the set of 841 vertebrates non-redundant position frequency matrices. The Matrix-Scan tool from RSAT^67^ was applied to scan the sequences, requiring a maximum p-value per site of 0.1. Background frequencies were calculated from the input set of sequences (markov=1, bg_pseudo=0.01). This scan resulted in a total of 5,389,949 motif hits across the 641 promoter sequences. Hits were further filtered on their sequence-matching score, so as to keep for each transcription factor hits above the median hit-score, resulting in a total subset of 453,101 motif hits. Finally, for each transcription factor, a Fisher test was applied to assess whether that TF exhibited enriched binding sites (relative to other TFs) within the promoter sequences of the up-regulated miRNAs (N=14), compared to a background set of promoter sequences corresponding to all other expressed miRNAs (N=91) A total of 215 TFBS motifs presented an odds-ratio greater than 1, indicating a relatively greater proportion of candidate promoters with a motif hit over other promoters. Filtering for odds-ratio greater than 2 and raw P-value lower than 0.1 yielded the set of 16 transcription factors whose binding sites are enriched among the concordant up-regulated miRNAs and that therefore represent potential regulators of dormancy.

### CUT&Tag

#### Sample preparation

Frozen nuclei were used and CUT&Tag was performed as described previously^68^ with minor modifications. Cells were dissociated using accutase and washed in PBS. Nuclei were extracted by resuspending the cells in ice-cold NE1 buffer (20 mM HEPES-KOH, pH7.5, 10 mM KCl, 0.5 mM Spermidine, 0.1% Triton X-100, 20% glycerol, 1 mM PMSF, 5 mM NaF, 1 mM Na3VO4) and incubating on ice for 10 min. Nuclei were fixed by incubating with 0.1% formaldehyde at room temperature for 2 min. 1.25 M glycine was added to stop cross-linking. Fixed nuclei were centrifuged for 4 minutes at 1300g at 4°C and were frozen in Wash buffer (20 mM HEPES-KOH, pH 7.5, 150 mM NaCl, 0.5 mM Spermidine, 1 mM PMSF, 5 mM NaF, 1 mM Na3VO4) with 10% DMSO at -80°C until further processing. For each CUT&Tag reaction, 3.5 µl of Concanavalin A beads (Bangs Laboratories) were equilibrated by washing 2 times in 100 µl Binding buffer (20 mM HEPES-KOH, pH 7.5, 10 mM KCl, 1 mM CaCl2, 1 mM MnCl2) and concentrated again in 3.5 µl Binding buffer. 10,000 cryopreserved nuclei were thawed and bound to the Concanavalin A beads by incubating for 10 min with rotation at room temperature. The beads were separated on a magnet and resuspended in 25 µl of cold Antibody buffer (Wash buffer with 0.1% BSA) containing respective primary antibody or IgG (TFE3, Sigma HPA023881, 1:50; IgG, Abcam ab46540, 1:100). Primary antibody incubation was done by incubating the beads at 4 °C with gentle nutation overnight. The beads were separated from the primary antibody on a magnet, resuspended in 25 µl of Wash buffer containing matching secondary antibody (guinea pig α-rabbit antibody, ABIN101961, Antibodies online, 1:100) and were incubated for 30 min at room temperature with gentle nutation. Afterwards, the beads were separated from the secondary antibody and washed once in 200µl of Wash buffer while being kept on the magnet. Homemade 3xFLAG-pA-Tn5, preloaded with Mosaic-end adapters, was diluted 1:250 in wash buffer and 25 µl of it was used to resuspend the beads. The beads were incubated for 1h at room temperature with gentle nutation and were washed once in 200 µl wash buffer. Tagmentation was performed by incubating the beads in 50 µl Tagmentation buffer (10mM MgCl2 in wash buffer) for 1h at 37°C. Tagmentation was stopped by adding 2.25 µl of 500 mM EDTA and 2.75 µl of 10% SDS to the beads followed by adding 0.5 µl of Proteinase K (20 mg/ml). After vortexing for 5 sec, the beads were incubated at 55 °C for 1h to solubilize the DNA fragments and then at 70 °C for 30 min to inactivate the Proteinase K. The beads were removed on the magnet and the DNA was purified using Chimmun DNA Clean & Concentrator (D5205, Zymo Research) following the manufacturer’s manual. The final CUT&Tag DNA was eluted in 25 µl of elution buffer. To amplify the NGS libraries, 25 µl of NEBNext HiFi 2x PCR Master Mix (New England BioLabs) was mixed with 21 µl of the CUT&Tag DNA, 2µl of 10 µM i5-and 2µl of 10 µM i7-unique barcoded primers^69^ and the following program was run on a thermocycler: 72 °C for 5 min, 98 °C for 30 sec, 98 °C for 10 sec, 63 °C for 10 sec (14 cycles of step 3-4) and 72 °C for 1 min. Ampure XP beads (Beckman Coulter) were used for post-PCR cleanup. 55 µl (1.1x volume) of the Ampure XP beads were added to the PCR mix and incubated for 10 minutes at room temperature. The beads were separated and washed two times using 80% ethanol on a magnet. Finally, the CUT&Tag libraries were eluted in 25 µl of Tris-HCl, pH 8.0. The quality of the CUT&Tag libraries was assessed by Agilent High Sensitivity D5000 ScreenTape System and QubitTM dsDNA HS Assay (Invitrogen). Sequencing libraries were pooled in equimolar ratios. The libraries were sequenced on Illumina next next-generation sequencer in paired-end mode at 5-8 million fragments per library.

#### Analysis

Raw reads were subjected to adapter and quality trimming with cutadapt (version 2.4; parameters: --quality-cutoff 20 --overlap 5 --minimum-length 25 --adapter AGA TCGGAAGAGC -A AGA TCGGAAGAGC) as were their respective input samples. Reads were aligned to the mouse genome (mm10) using BWA with the ‘mem’ command (version 0.7.17, default parameters). A sorted BAM file was obtained and indexed using samtools with the ‘sort’ and ‘index’ commands (version 1.10). Duplicate reads were identified and removed using GATK (version 4.1.4.1) ‘MarkDuplicates’ and default parameters. After careful inspection and validation of high correlation, replicates of treatment and input samples were merged respectively using samtools ‘merge’.

The smoothed Counts Per Million (CPM) signal was aggregated over extended promoter regions (upstream : 1,500 nt, downstream : 500 nt) by taking the maximum value over the regions and scaling those by the 95th quantile value across promoters for each condition. To further quantify the increased or decreased signal in paused over control samples, the log2 ratio of these maximum values was calculated, yielding for each promoter a positive value for relative increase signal, or a negative value for relative decrease. Finally, a one-sided Mann-Withney U test was performed to evaluate whether promoters of candidate miRNAs presented a greater increase in signal in paused samples, as compared to promoters of other measured miRNAs with available signal (N=88 from the previous 91 promoters).

### Nascent RNA expression analysis

25,000 wild-type or D*gcr8* KO cells in normal or mTORi conditions were plated onto 0.1% gelatin-coated glass coverslips. EU staining was performed using Click-iT RNA imaging kit (Invitrogen, C10329) following the manufacturer’s protocol. The coverslips were then mounted with Vectashield (Vector labs, H-1000) and sealed with nail polish. Imaging was done using a Zeiss LSM880 Airy microscope using Airy scan mode and image processing was done using Zen black and Zen Blue software (version 2.3). Image quantification was done using CellProfiler (version 4.2.1) where nuclei that were denoted as primary objects were identified and the normalized intensities of the EU staining were measured against the nuclear area (https://cellprofiler.org/). Data were plotted using GraphPad Prism (version 9).

### Differential gene expression analysis

Transcript abundance estimations per sample were obtained with kallisto^70^ and imported to R with the Bioconductor package tximport^71^. tximport summarizes transcript level abundance estimates for gene-level analysis. Differential gene expression was performed with the Bioconductor package DESeq2. Differentially expressed genes between WT paused vs normal and *Dgcr8* KO paused vs normal was defined as those with an adjusted p-value of 0.05 or smaller and |log2FC|>1.

### Differential isoform usage analysis

Abundance estimates per sample were obtained with kallisto. The R Bioconductor package IsoformSwitchAnalyzeR^72^ was used for differential isoform usage (DIU) analysis. Identification of differentially used isoforms across all genes with IsoformSwitchAnalyzeR is done through DEXseq^73^, which is a statistical method originally developed for differential exon usage based on the likelihood ratio test that has since been shown to adequately control for false discovery rate (FDR) in the setting of DIU.

### Sashimi plots

Reads were aligned to the mouse genome (build GRCm38/mm10) using STAR^74^ with default settings. Sashimi plots were done using ggsashimi^75^. Only differentially used transcripts were used for plotting.

### Western Blot (WB)

Cells were lysed with RIPA buffer supplemented with protease inhibitor cocktail for 30 min at 4C. The cells were spun at 14000 rcf for 20 min at 4C and the supernatant was collected for BCA protein quantification. 50 µg of protein was loaded onto a gradient gel (Biorad, 4561083) and the run was performed at 75V for 2.5h. Proteins were transferred to PVDF membrane (Invitrogen, IB24002) and blocking was done with 5% non-fat milk for 1h at room temperature. MBD2 or GAPDH primary antibodies were diluted in non-fat milk (1:1000) and incubated at 4C overnight. The blots were washed thrice in PBST for 10 minutes. The blots were then incubated with secondary antibody diluted 1:2000 and incubated for 1h at room temperature. The blots were washed thrice in PBST for 10 minutes each. Protein detection was done using SuperSignal West Dura on Biorad ChemiDoc MP Imaging System.

## SUPPLEMENTARY TABLES

**Table S1** Raw counts of wild-type and *Dgcr8* KO ESCs in normal and pausing conditions, and results from differential gene expression analysis

**Table S2** Full list of GO terms

**Table S3** List of identified small RNAs and expression levels in all conditions

**Table S4** Relative expression levels of identified miRNAs

**Table S5** Nanostring output

**Table S6** List of identified proteins and expression levels in normal and paused mouse ESCs

**Table S7** miRNA-target networks node and edge quantifications

